# Multimodal Magnetic Resonance Imaging-based neuromarkers trace longitudinal changes in cognitive functioning in Attention Deficit/Hyperactivity Disorder

**DOI:** 10.64898/2026.03.07.710313

**Authors:** K. Jack Scott, Kseniia Konopkina, Farzane Lal Khakpoor, Irina Buianova, William van der Vliet, Narun Pat

**Author notes:** All correspondence to: Jack Scott Department of Psychology, University of Otago, William James Building, 275 Leith Walk, Dunedin 9016, New Zealand.

## Abstract

The National Institute of Mental Health’s Research Domain Criteria (RDoC) framework conceptualises cognition as a core functional domain for psychopathology that should be studied across multiple units of analysis and developmental timescales. In ADHD, however, it remains unclear whether neuroimaging-derived markers can not only predict inter-individual differences in cognition but also track longitudinal cognitive development and capture cognition–psychopathology relationships. Using the longitudinal Oregon ADHD-1000 study (n = 594 participants; 1,053 observations), we developed multimodal machine-learning markers of general cognitive functioning (g) from structural and resting-state functional MRI. The multimodal marker achieved an out-of-sample correlation of *r* = .46 and generalised similarly across children with and without ADHD. The marker explained 25.01% of interindividual variance (*r* = .48) and 18.82% of intraindividual variance (*r* = .52) in cognitive functioning. Commonality analyses showed that it captured 60.87% of intraindividual age-related cognitive variation (5.18% of the total variance in g). The marker also accounted for substantial portions of the cognition–hyperactivity association (58.79%; 6.39% of total variance in g) and the cognition–inattention association (25.99%; 4.13% of total variance in g). These findings provide evidence that multimodal structural and functional MRI can generate RDoC-informed markers that predict cognitive functioning, track cognitive development over time, and capture meaningful links between cognition and ADHD symptoms. Although predictive performance remains below levels required for clinical application, the results establish a foundation for longitudinal, RDoC-inspired investigations of cognitive development in ADHD.

## Introduction

Attention Deficit/Hyperactivity Disorder (ADHD) affects about 7–8% of children worldwide and often persists into adulthood^1–3^, impacting academic performance^4^, occupational success^5,6^, and quality of life^7^. Individuals with ADHD have an all-cause mortality rate nearly twice that of neurotypical peers^8^. The disorder is associated with difficulties across several cognitive domains, including executive functioning, working memory, processing speed, and sustained attention^9–12^, which are strongly linked to long-term functional outcomes^4–6,13^.

One influential framework for investigating cognitive functioning in ADHD is the National Institute of Mental Health’s Research Domain Criteria (RDoC) ^14,15^. Rather than conceptualising mental disorders as discrete diagnostic categories, the RdoC framework emphasises the study of functional domains, not the disorder such as ADHD per se, that span a continuum from typical to atypical behaviour. Cognitive systems constitute one of the framework’s core domains, alongside negative valence systems, positive valence systems, social processes, arousal and regulatory systems, and sensorimotor systems. Consistent with this approach, for instance, Karalunas and colleagues demonstrated that some cognitive aspects (e.g., working memory performance) predicted symptom remission in individuals with ADHD ^16^, highlighting the value of dimensional cognitive measures for understanding clinical outcomes.

The RDoC framework extends beyond behavioural assessments of cognition by conceptualising cognitive functioning across multiple neurobiological units of analysis, including neural systems, physiology, molecular processes, and genetics^17^. This multilevel perspective encourages investigation of the biological mechanisms underlying cognitive functioning and their relationship to psychopathology^14,18,19^. In ADHD research, structural and functional magnetic resonance imaging (sMRI and fMRI) are among the most widely used tools for examining neural differences between individuals with and without ADHD, traditionally through case–control comparisons and symptom correlations^20,21^. Common sMRI measures include cortical thickness, surface area, and subcortical volume, whereas resting-state fMRI most often focuses on functional connectivity, defined as the temporal correlation of BOLD signals between brain regions over the course of a scan. Despite the increasing use of neuroimaging in ADHD research, only a few studies have examined the extent to which sMRI and fMRI measures can predict inter-individual differences in cognitive functioning across individuals with and without ADHD^18,22^.

In population-based samples, machine-learning approaches have successfully predicted inter-individual differences in cognitive performance from sMRI and fMRI data^23–25^. More recently, these methods have been extended to ADHD cohorts, showing promising performance in predicting cognitive functioning from structural and resting-state neuroimaging measures^18,22^. For example, Chopra and colleagues developed machine-learning models using resting-state fMRI data from large population-based samples and demonstrated that the resulting models could successfully predict cognitive functioning across several psychiatric conditions, including ADHD^18^. Similarly, Khakpoor and colleagues integrated structural and functional neuroimaging features using a multimodal stacking framework to predict cognitive functioning^22^. Their results showed that combining information across MRI modalities improved predictive performance relative to single-modality models and generalised well to children with ADHD. More importantly, they also showed the generalisability of their predictive models built from large population-based data to a smaller, case-control for ADHD dataset. Together, these findings suggest that machine-learning approaches can train neuroimaging markers that capture meaningful inter-individual differences in cognitive functioning across both ADHD and non-ADHD populations.

Still so far, resting-state fMRI studies have predominantly focused on functional connectivity, while other informative metrics, such as Regional Homogeneity (ReHo)^26^ and Amplitude of Low-Frequency Fluctuations (ALFF)^27^, have received comparatively little attention. ReHo quantifies the local synchronisation of BOLD signals across neighbouring voxels or vertices^26^, whereas ALFF measures the magnitude of spontaneous neural activity by calculating the power of low-frequency BOLD fluctuations^27^. Both ReHo and ALFF have been shown to differ in individuals with ADHD^28,29^ and can contribute to diagnostic prediction^30^. Moreover, evidence indicates that integrating multiple neuroimaging modalities and feature sets through machine-learning approaches such as multimodal stacking can improve predictive performance^31–33^. Nevertheless, it remains unclear whether combining a broader range of sMRI and resting-state fMRI features, including ReHo and ALFF, can enhance the prediction of cognitive functioning across individuals with and without ADHD.

Beyond examining inter-individual differences in cognitive functioning, the RDoC framework emphasises the developmental nature of functional domains^15^. Consequently, for fMRI and sMRI measures to qualify as RDoC-informed markers of cognitive functioning in ADHD research, they must not only capture between-person variation in cognition but also sensitively track within-person changes over time^32,34^. This requirement presents a challenge, as resting-state fMRI measures have generally shown low test–retest reliability, raising concerns about their suitability for detecting longitudinal changes in cognitive functioning ^35^. However, recent evidence suggests that machine-learning approaches that aggregate information across the full connectome can substantially improve the reliability of fMRI-derived markers ^32,33,36^. Furthermore, Tetereva and colleagues demonstrated that integrating fMRI and sMRI data through a multimodal stacking framework enhances both predictive performance and test–retest reliability. Together, these findings indicate that machine-learning approaches, particularly multimodal stacking, may enable s/fMRI-derived markers to track longitudinal changes in cognitive functioning in individuals with and without ADHD. The extent to which these markers capture within-person change can be evaluated in longitudinal studies using methods such as linear mixed-effects variance decomposition^37^.

However, demonstrating sensitivity to within-person change is not, by itself, sufficient to establish that an s/fMRI-based marker captures the developmental nature of an RDoC functional domain. A marker may track changes in an individual’s state without necessarily indexing developmental processes. For example, EEG can reliably track fluctuations in sleep or arousal, yet its sensitivity to these short-term states does not, in itself, demonstrate relevance to brain maturation over years. Thus, the ability to predict within-person cognitive change must be complemented by evidence that the marker reflects year-long developmental processes underlying cognitive growth.

Cognitive functioning generally improves with age, including among children with ADHD in parallel with ongoing brain maturation^38–41^. An important question, therefore, is the extent to which s/fMRI-based markers account for the association between cognitive functioning and age. Addressing this question helps determine whether these markers capture neurodevelopmental variation that is meaningfully related to cognitive development, rather than nonspecific fluctuations in brain state. In longitudinal data, this contribution can be quantified using commonality analysis, which partitions variance in cognitive functioning into components uniquely and jointly explained by age and neuroimaging-based markers^42,43^. Such analyses provide a direct test of whether neuroimaging markers account for the developmental coupling between cognition and age that is central to the RDoC framework.

Finally, establishing the clinical relevance of RDoC-inspired neuroimaging markers of cognition for ADHD requires demonstrating that these markers capture the association between cognitive functioning and ADHD symptomatology—the cognition– psychopathology relationship central to the RDoC framework^17^. ADHD is characterised by two core symptom dimensions: a) inattention, defined by difficulties sustaining attention, maintaining focus, and organising goal-directed behaviour, and b) hyperactivity, characterised by excessive motor activity, restlessness, and difficulties with behavioural inhibition^44^. Cognitive functioning has been often linked to both symptom dimensions^45–48^. Consequently, clinically meaningful neuroimaging markers of cognition should not only predict cognitive functioning but also capture its association with ADHD symptoms at both the between-person and within-person levels. Following this rationale, commonality analysis can be used to quantify the variance in cognitive functioning that is uniquely and jointly attributable to neuroimaging-based markers and ADHD symptom dimensions, thereby evaluating the extent to which these markers capture the cognition–psychopathology relationship emphasised by the RdoC framework^42,43^.

In the current study, using large-scale longitudinal sMRI and fMRI data from the Oregon-1000 ADHD Study^49^, we trained machine-learning models to predict cognitive functioning, operationalized as the general cognitive factor (g), in children and adolescents with and without ADHD (see Figure 1 for the overall study design). The models incorporated four structural MRI feature sets (cortical surface area, cortical thickness, subcortical volume, and total brain volume) and three resting-state functional MRI feature sets (functional connectivity, ALFF, and ReHo)^50^. Predictions from these modalities were subsequently integrated using a multimodal stacking approach.

**Fig. 1:**
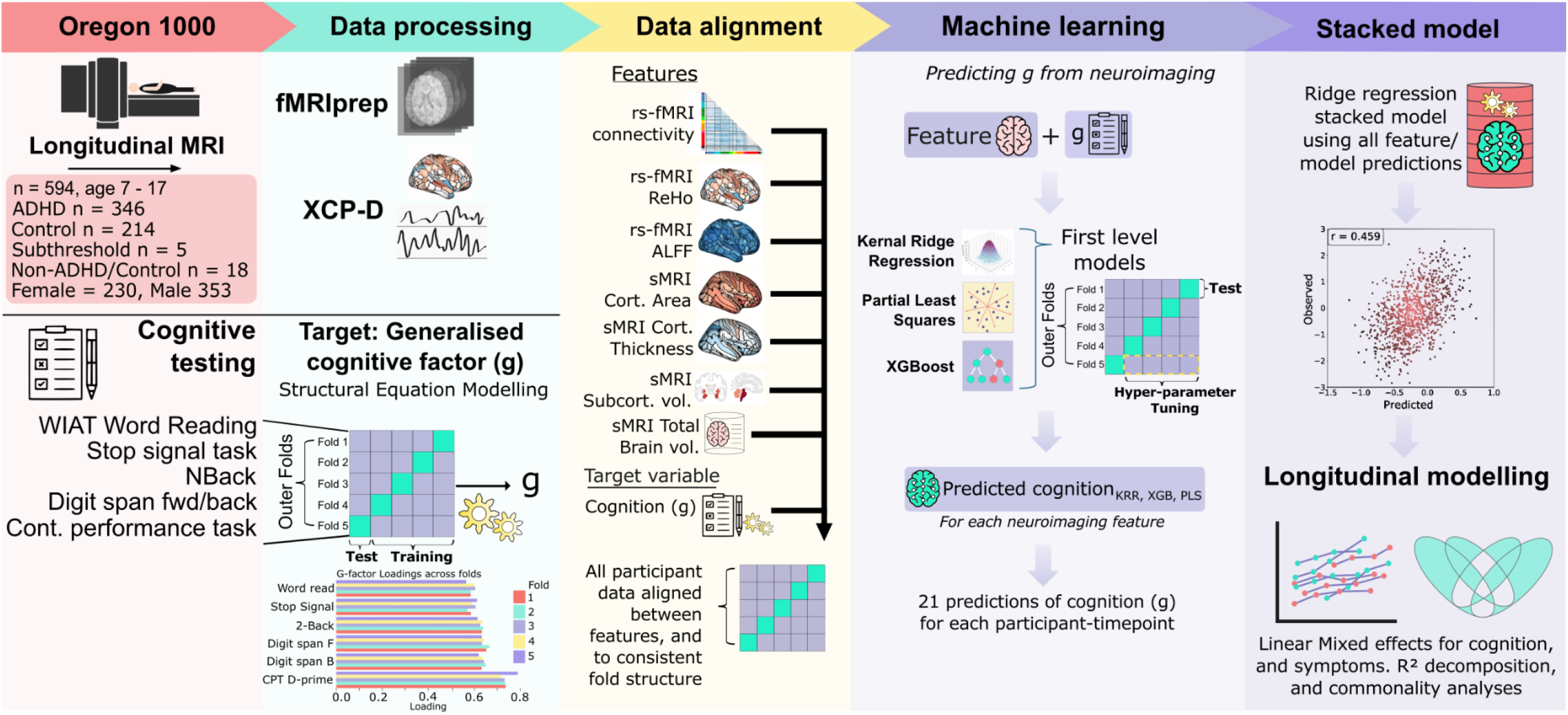
Study overview. The Oregon-1000 ADHD longitudinal dataset (final composition following processing: n=594, ages 7-17, 1,053 timepoints) contained structural and functional MRI alongside cognitive testing. Following preprocessing, seven neuroimaging features were extracted: resting-state fMRI (functional connectivity, ALFF, ReHo) and structural MRI (cortical surface area, cortical thickness, subcortical volume, total brain volume). To derive a general cognitive factor (g), we applied confirmatory factor analysis to six cognitive test scores. Three machine learning algorithms (Kernel Ridge Regression, Partial Least Squares, XGBoost) were trained on each feature separately using nested cross-validation, producing 21 first-stage predictions. A ridge regression stacked model combined these predictions to generate final cognitive estimates. Linear mixed-effects models decomposed predictions into intraindividual trajectories and interindividual differences, with commonality analyses examining relationships between brain predictions, age, and ADHD symptoms.

Guided by the RDoC framework^15^, this study had four overarching aims. First, we tested whether combining a broader range of structural and resting-state functional neuroimaging features, including ALFF and ReHo, through multimodal stacking would improve the prediction of cognitive functioning across individuals with and without ADHD. Second, using linear mixed-effects variance decomposition^37^, we quantified the extent to which the resulting multimodal neuroimaging markers explained intra-individual (within-person) and inter-individual (between-person) variation in cognitive functioning. Third, using linear mixed-effects commonality analysis^42,43^, we examined the degree to which these multimodal neuroimaging markers accounted for the association between cognitive functioning and age. Finally, again using commonality analysis^42,43^, we investigated the extent to which the multimodal neuroimaging markers captured the relationship between cognitive functioning and ADHD symptoms.

## Materials and Methods

### Dataset and participants

The Oregon-1000 ADHD study^49^, an accelerated (lag) longitudinal dataset, provided cognitive, neuroimaging and ADHD rating-scale data^44^. At baseline, individuals were excluded if they used non-stimulant psychotropic medications, had a history of epilepsy or seizures, an IQ below 80, or specific comorbid mental health disorders (including current severe depressive episode, psychosis, tic disorder, head injury, autism spectrum, or other health condition). During follow-up, severe depressive episodes were not an exclusion criterion, as it was considered a trajectory outcome^49^. Participants on stimulant medication discontinued use 48 hours before cognitive or neuroimaging assessments^49^. Among ADHD participants, 35% were prescribed stimulants: 36.4% amphetamine formulations, 51.3% methylphenidate, and 14.3% other stimulants.

The sample was predominantly White (n = 452, 76.2%), followed by individuals identifying as more than one race (n = 60, 10.1%), Black or African American (n = 9, 1.5%), Asian (n = 9, 1.5%), American Indian or Alaska Native (n = 1, 0.2%), and race not specified (n = 64, 10.8%). With respect to ethnicity, 32 participants (5.4%) identified as Hispanic or Latino, 499 (84.0%) identified as not Hispanic or Latino, and ethnicity was not specified for 63 participants (10.6%). Demographic characteristics are presented in Supplementary Table S1.

ADHD symptom severity was assessed using the ADHD Rating Scale-IV^44^, administered at the time of neuroimaging or within six months of the scan (see Supplementary Table S2 for data coverage). Raw scores for the inattention and hyperactivity/impulsivity subscales were used in all analyses. Raw scores were selected because they are not age-standardized, allowing symptom trajectories to be modelled directly using linear mixed-effects analyses. Symptom ratings were obtained from participants’ parents and were independently evaluated by two clinicians^49^.

The study provided neuroimaging data, along with associated cognitive, psychopathology, and demographic measures, from 644 participants. Following data postprocessing and the alignment of cognitive, neuroimaging, and ADHD rating-scale data based on matched functional connectivity matrices, the final analytic sample consisted of 594 participants, ages 7-17 years old (*M* age at baseline = 10.2 years, *SD* = 1.55 years), including 230 females (38.7%), 353 males (59.4%), and 11 for whom assigned sex was absent (1.9%). Across participants, 1,053 observations (time points) were available, with individuals contributing between 1 and 5 time points (M = 1.77, SD = 0.96). Specifically, 311 participants contributed one timepoint, 150 contributed two, 92 contributed three, 39 contributed four, and 2 contributed five. Consecutive assessments were separated by an average interval of 23 months (SD = 15.8 months).

At each study visit, clinicians in the Oregon ADHD-1000 study assigned participants to one of five phenotypic categories based on clinical assessment: Control (no elevated ADHD symptoms), ADHD (met diagnostic criteria for ADHD), Subthreshold (ADHD symptoms exceeded normative levels for typically developing controls but did not meet diagnostic criteria), Neither Clean Control nor ADHD (participants who initiated treatment with other psychotropic medications or received additional diagnoses beyond ADHD after enrolment), and Not Assessed (no phenotype rating available for that visit).

Because phenotypic classification could vary across visits, a stable phenotype was derived for each participant. Participants who received an ADHD classification at any visit were assigned to the ADHD group. Among the remaining participants, those classified as Control more frequently than Neither Clean Control nor ADHD were assigned to the Control group, whereas those classified as Neither Clean Control nor ADHD more frequently, or with equal frequency, were assigned to that group. Visits coded as Not Assessed were excluded from these comparisons, and classification was based on available non-missing ratings. Participants were classified as Not Assessed only if no valid phenotype rating was available across all visits.

Applying these criteria, the final sample included 214 Control participants, 346 ADHD participants, 5 Subthreshold participants, 18 Neither Clean Control nor ADHD participants, and 11 Not Assessed participants.

All 594 participants were included in the machine-learning analyses predicting cognitive functioning from s/fMRI data. However, longitudinal analyses using linear mixed-effects models were restricted to participants with at least two neuroimaging assessments, including 107 females (37.81%), 174 males (61.48%) and 2 participants with sex not available. The mean age of this subsample was 11.0 years (*SD* = 2.0, range = 7.2–16.6 years). Among participants classified as having ADHD, 104 were female (30.1%) and 242 were male (69.9%), with a mean age of 10.8 years (*SD* = 1.7, range = 7.3–15.6 years).

No participants were excluded on the basis of phenotype assignment. Thus, all 594 participants who passed quality-control procedures were retained for the machine-learning analyses. Participants were excluded only from the longitudinal analyses when multiple neuroimaging time points were required.

### Target of machine-learning prediction: Cognitive functioning

We analysed six cognitive measures representing several key cognitive constructs: N-Back 2-back accuracy^51^ (working memory and attention), Continuous Performance Task (CPT) d’ catch trials^52^ (sustained attention and vigilance; scored as z(Hit rate) – z(False alarm rate)), Stop Signal Reaction Time Task (SSRT) reaction time standard deviation^53^ (inhibitory control), and Digit Span Forward (DSF), and Backward (DSB) tasks (working memory; scored as longest correctly recalled sequence) from the Wechsler Intelligence Scale for Children IV^54^, and Wechsler Individual Achievement Test (WIAT) word reading score^55^ (phonological awareness and decoding). We reverse-coded the stop signal reaction time standard deviation scores to match the directionality of all other scores, and also shifted the log-transformed CPT d’ catch to improve normality and handle negative values.

Data coverage across cognitive measures averaged 95.92% (Table S2). Variables were selected based on high longitudinal coverage, temporal proximity to neuroimaging assessments (within six months of each scan), and suitability for latent variable modelling, as evaluated through intercorrelations (Table S3).

To derive a general cognitive ability factor (g), we conducted a confirmatory factor analysis (CFA)^56^ using a first-order model in which all cognitive measures loaded directly onto a single latent g factor (Figure 1, Table S4). The CFA dataset comprised 1,278 observations across all phenotype groups. Cognitive measures were iteratively refined to achieve stable model convergence. This process resulted in the exclusion of WIAT Mathematical Reasoning, SSRT mean reaction time, SSRT Go accuracy, SSRT probability of stopping, and CPT d′ stimulus. The latter was removed because it was highly correlated with CPT d′ catch trials (*r* = .85), indicating substantial redundancy.

The factor score of g was extracted using the lavPredict function in the lavaan package, which applies the regression method by default. The g score served as the target variable in subsequent machine learning analyses. To prevent data leakage, CFA models were fitted separately within each training set across the five outer cross-validation folds, and the corresponding fitted models were then applied to the held-out test sets. Figure 1 presents the task loadings obtained across folds.

We selected a first-order model because the cognitive measures were drawn from different assessment batteries, and there was no strong a priori theoretical basis for specifying a more complex cognitive structure. Moreover, our primary interest was the extracted g scores rather than the latent structure of the cognitive measures themselves. Nevertheless, to evaluate the robustness of our findings to alternative specifications of cognitive structure, we conducted a sensitivity analysis using the full dataset. Specifically, we compared g scores derived from the first-order CFA model with those obtained from a second-order CFA model. The second-order model included the same six cognitive measures, which loaded onto two first-order factors that, in turn, loaded onto a higher-order g factor. The two first-order factors were identified based on exploratory factor analysis (see Supplementary Materials). All analyses were conducted in R (v4.5.1) using the lavaan (v0.6-19) and psych (v2.5.6) packages.

### Features used for machine-learning prediction: s/fMRI data

Participants were scanned at Oregon Health & Science University’s Advanced Imaging Research Center using a 3.0 Tesla Siemens Tim Trio Magnetom MRI scanner, equipped with a 12-channel head coil. The authors reported no changes to this equipment or acquisition/software throughout the study period^49^. For full MRI scanning parameters, refer to the original article, specifically the Supplementary Materials^49^. We received a minimally pre-processed BIDS version of this dataset and pre-processed the data as described in the following two sections. We applied framewise displacement-based quality control, excluding runs (each comprising five minutes of fMRI data) in which more than 50% of volumes exceeded a 0.5 mm threshold. A minimum of two runs passing this criterion were required for a participant-time point to be retained. Participant-timepoints failing this threshold were excluded across all modalities, including structural MRI (101 excluded timepoints), so as to ensure multimodal data completeness at each retained timepoint

### Structural MRI (sMRI)

sMRI data consisted of T1-weighted anatomical images acquired with the following parameters: TR = 2300 ms, TE = 3.58 ms, sagittal orientation, 256 × 256 matrix, and slice thickness = 1 × 1 × 1.1 mm. Four sMRI feature sets were derived using Freesurfer (7.3.2), executed as part of fMRIPrep (24.0.0)^57^: cortical surface area, cortical thickness, subcortical volume, and total brain volume. The first three feature sets were parcellated using the Destrieux atlas (148 cortical regions)^58^ and FreeSurfer automatically segmented brain volume (ASEG) atlas (19 subcortical regions)^59^, resulting in 167 features per set. The total brain volume feature set comprised five global features: intracranial volume, total cortical gray matter volume, subcortical gray matter volume, total white matter volume, and the ratio of brain segmentation volume to estimated intracranial volume. Specifically, the total brain volume is a global measure representing the aggregate volume of the entire brain, as opposed to regional sMRI metrics in the case of cortical surface area, cortical thickness, subcortical volume. See Supplementary materials for additional details on sMRI data processing.

### Functional MRI (fMRI)

fMRI data consisted of three 5-minute runs of resting-state BOLD data, during which participants fixated on a white cross against a black background ^49^. fMRI acquisition parameters included TR = 2500 ms, TE = 30 ms, flip angle = 90°, and voxel size = 3.75 × 3.75 × 3.80 mm. Our pipeline consisted of preprocessing using fMRIPrep (24.0.0)^57^, and post-processing using XCP-D (0.10.6) ^60^. XCP-D featured the aCompCor^61^ noise regression pipeline (up to 5 white matter, 5 CSF components; 12 motion parameters including 6 head motion parameters, their temporal derivatives all previously generated by fMRIPrep).

Three resting-state fMRI sets of features were derived from XCP-D, Functional Connectivity, Regional Homogeneity (ReHo^26^) and Amplitude of Low-Frequency Fluctuations (ALFF ^27^). All of these fMRI sets of features were parcellated during XCP-D processing, using the Glasser atlas (360 cortical regions)^62^ and FreeSurfer ASEG atlas (19 subcortical regions)^59^, which we then concatenated. Note that we excluded two cortical regions, the left and right hippocampal parcels from the Glasser atlas, due to poor coverage in many participants, resulting in a total of 377 regions.

For functional connectivity, we computed the Pearson correlation (r) between censored time series (FD < 0.5mm) of each pair of regions, yielding 70,876 unique connections, and applied Fisher’s z-transformation. For ReHo, we used the same censored time series and assessed local similarity using a 27-voxel neighbourhood (including face, edge, and corner neighbours) and Kendall’s coefficient of concordance ^60^, calculating mean ReHo for each parcellated region, yielding 377 features for ReHo. For ALFF, specifically mean ALFF (mALFF), we calculated the power spectral density from uncensored time series, band-pass filtered at 0.01–0.1 Hz, and standardised by the global mean ALFF. The ALFF was then averaged for each parcellated region, resulting in 377 features for ALFF. See Supplementary materials for additional details on fMRI data processing.

### Machine-learning processes

To predict participants’ cognitive functioning from s/fMRI features, we used a two-stage stacking procedure. In the first stage, machine learning algorithms were trained to predict cognitive functioning (*g* score) from one neuroimaging feature set (e.g. cortical surface area). In the second “stacked-modelling” stage, we combined the first-stage predictions from each combination of an algorithm and feature sets and used these predictions as input to predict *g*. To prevent data leakage, participant data were entirely contained within either training or test sets, and never split across folds. Performance was assessed using Pearson r, Coefficient of Determination (R^2^), Mean Absolute Error (MAE), and Mean Square Error (MSE).

### First stage models

Following alignment of g-scores and participant neuroimaging data to the same fold structure, we trained three functionally distinct machine-learning algorithms, seeking to use these algorithms in a complimentary fashion to capture the brain-cognition relationship. Kernel Ridge Regression (KRR) provides regularized linear modelling that handles multicollinearity well, with optional kernel transformations for non-linear relationships^63,64^. Partial Least Squares (PLS) regression performs dimensionality reduction by identifying latent components and maximising feature-outcome covariance^65,66^. XGBoost uses gradient-boosted decision trees to capture complex non-linear interactions without a priori functional assumptions ^67^. Hyperparameter grids for each algorithm are provided in Supplementary Materials (Tables S5, S6).

We implemented 5-fold nested cross-validation (CV), with nested-CV utilised for hyperparameter tuning using grid search. For each outer fold (80% training, 20% test), we performed inner 5-fold cross-validation for hyperparameter tuning using the training set. Optimal hyperparameters were selected by minimizing MSE (XGBoost, PLS) or MAE (KRR), then models were retrained on the full outer training set and evaluated on the held-out test set. This produced 21 first-stage predictions (7 features × 3 algorithms) for each participant-timepoint.

### Stacked model

Following previous research^32,68–70^, we trained a stacked machine-learning model to combine the 21 first-stage predictions into a single prediction for each participant at each timepoint To prevent data leakage, we used the same nested cross-validation (CV) framework as in the first-stage models. Within each outer fold, a ridge regression model was trained exclusively on training-set predictions generated by the first-stage models, without any involvement of the test set. Thus, the stacked model learned from the in-sample predictions of g produced by each model–feature combination training set (Figure 1), rather than directly from the brain features themselves. Hyperparameter tuning was performed within the training data of each outer fold using an alpha grid ranging from 10^−2^ to 10^5^. The hyperparameter configuration yielding the highest R^2^ was selected, and the final ridge regression model was fitted using the auto solver with an intercept term.

To assess the association between each first-stage prediction and the stacked model, we computed Haufe feature-importance scores^71^. Raw ridge regression coefficients can be misleading when input features are correlated, as shared variance between features can suppress or inflate individual weights^71,72^. Haufe feature-importance addresses this by transforming model weights into activation patterns, reflecting the correlation between each input and the model’s predicted values, and is thus robust to multicollinearity among first-stage predictions.

### Aim#1: Comparing stacked vs. first-stage models

We conducted bootstrap analyses to evaluate the predictive performance of stacked model, as compared to that of the first-stage model. We focused this analysis on the observed and predicted g-scores pools across all outer-fold test sets. At each of 5,000 bootstrapping iterations, we calculated Pearson-r correlations between observed and predicted g-scores for both the stacked model and each first-stage model. Correlations were Fisher z-transformed to normalize distributions. We then computed differences in these transformed correlations between stacked and first-stage models (z-stacked minus z-first-stage). Statistical significance was determined by whether the 95% confidence interval of the difference distribution excluded zero. Fisher z differences served as effect sizes, with larger absolute values indicating greater performance differences.

### ALFF and ReHo feature importance

To assess whether inclusion of ALFF and ReHo resulted in improved predictive performance, we trained an ablated stacked model that excluded ALFF- and ReHo-based predictions and instead relied solely on predictions derived from functional connectivity and structural feature sets. We then conducted a bootstrap analysis analogous to that used in the primary model evaluation, quantifying the difference in predictive performance between the ablated and full stacked models. Specifically, we computed the difference in Fisher z-transformed correlations across bootstrap samples to determine whether inclusion of ALFF and ReHo yielded a significant improvement in prediction accuracy.

### Linear mixed-effects models

We applied linear mixed-effects (LME) models, estimated using maximum likelihood, to the observed and predicted g-scores pooled across all outer-fold test sets. These analyses were restricted to the 283 participants who had more than one time point (742 time points in total), enabling the estimation of intraindividual cognitive changes.

In these LME models, observed g-scores (denoted g_observed_) were extracted directly from the cognitive measures via the first-order CFA, whereas predicted g-scores were derived from s/fMRI data using the stacked model. To distinguish intra-from inter-individual variability in the predicted g-scores, derived from s/fMRI, we computed two explanatory variables: ĝ_mean_ and ĝ_deviation_ ^73^. ĝ_mean_ was calculated as the participant-specific mean of predicted g-scores across all available timepoints and reflects interindividual variability. ĝ_deviation_ was computed as each timepoint’s deviation from that participant’s mean predicted g-score, capturing intraindividual variability in predicted g-scores over time. Both observed and predicted g-scores were z-standardised across the full sample.

Next, for LME models investigating age, we created age variables in a similar fashion to the predicted g-scores. We started by subtracting the participants’ minimum age (years) from their age at each time point. Then, we computed the mean age across all available timepoints, denoted age_mean_, and the deviation of each timepoint from that participant’s mean age, age_deviation_, to distinguish between inter- and intra-individual variability in age.

Similarly, for the models investigating ADHD Symptoms, including hyperactivity or inattention, we computed the mean of each symptom score across all available timepoints, denoted Hyperactivity_mean_ and Inattention_mean_, respectively, to reflect interindividual variability in symptom scores. Likewise, we computed each timepoint’s deviation from that participant’s mean symptom, denoted Hyperactivity_deviation_ and Inattention_deviation_, respectively, to reflect intraindividual variability in symptom scores. Hyperactivity and inattention scores were based on the ADHD rating scale IV^44^ and z-standardised across the full sample.

In all LME models, the observed g-scores served as the response variable. We varied the fixed effects of interest while including a participant-specific random intercept (grouped by participant ID) to account for repeated measurements. Although we initially considered random slopes, some models failed to converge when a random slope term was included. Because commonality analyses require the same random-effects structure across all models to ensure valid model comparisons, we adopted a random-intercept-only structure for all LMEs. This approach represented the most complex random-effects structure that was both estimable and consistent across the full set of models^74^.

The LME models were used to address four questions: (a) to test the generalisability of the predicted g-scores across ADHD diagnoses, (b) to decompose the extent to which predicted g-scores explained inter-versus intra-individual variance in observed g-scores, (c) to quantify the commonality between age and predicted g-scores, and (d) to quantify the commonality between ADHD symptoms and predicted g-scores.

### Generalisability of the predicted g-scores across ADHD diagnoses

Before examining how the neuroimaging marker captured variance in cognitive functioning, we first assessed whether its predictions generalised consistently to children with and without ADHD. To assess whether the stacked model predicted g-factor consistently across ADHD diagnoses, we fitted two LME models: an ADHD-additive model and an ADHD-interactive model.

Both ADHD-additive and ADHD-interactive models regressed observed g-scores on predicted g-scores and ADHD status. The ADHD-interactive model additionally included interaction terms between predicted g-scores and ADHD status, allowing the model to test whether the relationship between observed and predicted g-scores varied by diagnostic groups. If these interaction terms did not significantly improve model fit, we inferred that the stacked model predicted g-scores similarly across ADHD groups Model comparisons were based on the Likelihood Ratio Test, Bayesian Information Criterion (BIC), and changes in R^2^, with preference given to the most parsimonious model.

Because our primary focus was on ADHD diagnoses, this analysis was restricted to participants who either met clinical criteria for ADHD at one or more timepoints or were classified as controls for the majority of their assessed timepoints. Consequently, five participants were excluded from the additive versus interactive ADHD linear mixed-effects analyses because they had multiple neuroimaging timepoints without corresponding diagnostic data, or because the majority of their available assessments were classified by clinicians as neither ADHD nor control.

The ADHD-additive model included ĝ_mean_, ĝ_deviation_, and ADHD status (where 1=ADHD, 0=control) as fixed effects and incorporated a random intercept, using the following lme4 syntax:

#### ADHD Additive model

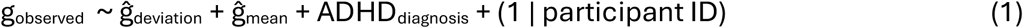

Similarly, the ADHD-interactive model added interaction terms between ĝ_mean_ and ĝ_deviation_ and ADHD status to the ADHD-additive model as fixed effects, using the following lme4 syntax:

#### ADHD Interactive model

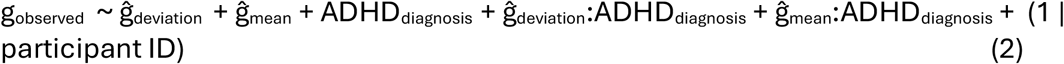

### Aim#2: Decomposition of intra-versus inter-individual variation

To assess how much of the inter-versus intra-individual variation in observed g-scores was accounted for by predicted g-scores, we performed variance decomposition on the neuroimaging-only model ^75^. In this model, we included both ĝ_mean_ and ĝ_deviation_ as fixed effects and incorporated a random intercept, using the following lme4 syntax:

#### Neuroimaging-only model

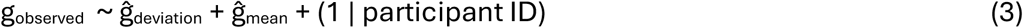

The total variance in observed g-scores was partitioned into level-1 and level-2 components. Because participant ID served as the grouping variable, the level-1 component reflects intraindividual (within-person) variation across sessions:

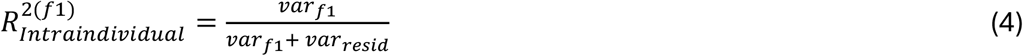

Here, 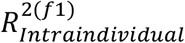 quantifies the proportion of intraindividual variance in observed g-scores explained by the level-1 fixed effect (i.e., ĝ_deviation_). The term *var*_*f*1_ denotes variance explained by this level-1 fixed effect, and *var*_*resid*_ is the level-1 residual variance.

The level-2 component reflects interindividual (between-person) variation across sessions:

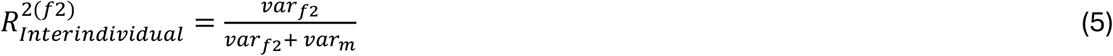

Here, 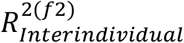 quantifies the proportion of interindividual variance in observed g-scores explained by the level-2 fixed effect (i.e., ĝ_mean_). The term *var*_*f*2_ denotes the variance explained by this level-2 fixed effect, and *var*_*m*_ corresponds to the random-intercept variance.

### Aim#3: Commonality between age and predicted g-scores

To quantify how well the stacked model captured age-related cognitive variation, we implemented two additional LME models. First, the age-only model served as a baseline, estimating the variance in observed g-scores explained solely by age. The age-only model included age_deviation_ and age_mean_ as fixed effects and had a random intercept to account for interindividual differences, using the following lme4 syntax:

#### Age-only model

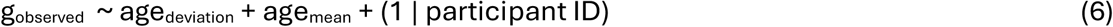

Second, the neuroimaging-and-age model enabled us to assess the extent to which age-related variation in cognitive functioning was accounted for by predicted g-scores using a commonality analysis^43^. This model included explanatory variables from both neuroimaging-only and age-only models, using the following lme4 syntax:

#### Neuroimaging-and-age model

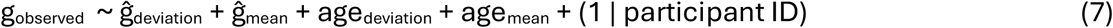

Commonality analysis^42,43^ partitions the total variance explained by a set of fixed-effect variables into components uniquely attributable to each fixed-effect variable and components shared among fixed-effect variables. We implemented this by computing marginal R^2^, the proportion of variance in observed g-scores explained by the fixed-effect variables, across all possible combinations of fixed-effect variables. Comparing marginal R^2^ values across these model combinations enabled us to quantify the unique and shared contributions of predicted g-scores and age to variance in observed g. When negative commonality coefficients arose, we followed a recommended practice^76^ by setting negative components to zero.

We defined age-related cognitive variation as the marginal R^2^ from a model in which age_deviation_ and age_mean_ were the only fixed effects. To determine how much of this age-related cognitive variation was explained by the stacked model, we computed the proportion between a) the commonality between the age variables and the predicted g-scores and b) the total variation due to each age variable.

### Aim#4: Commonality between ADHD symptoms and predicted g-scores

To examine how well the stacked model captured the relationship between cognition and ADHD symptoms, we fitted four additional LME models, two for each symptom dimension: hyperactivity and inattention.

As baseline models, each symptom model included the deviation and mean of each symptom score as fixed effects, and incorporated a random intercept. Here we test the relationship between each symptom and observed g-scores, using the following lme4 syntax:

#### Hyperactivity only model

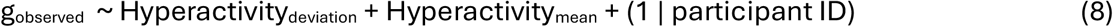

#### Inattention only model

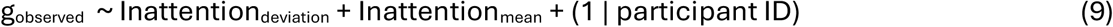

Similar to the neuroimaging-and-age model, including the deviation and mean of predicted g-scores for each symptom model enabled us to assess the extent to which symptom-related variation in cognitive functioning was accounted for by predicted g-scores using a commonality analysis^43^. We used the following lme4 syntax:

#### Neuroimaging-and-hyperactivity model

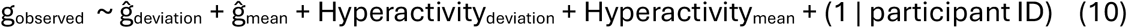

#### Neuroimaging-and-inattention model

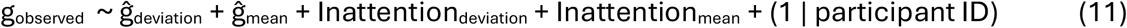

We applied commonality analysis to the symptom models using the same approach as for age-related variation in cognition. To quantify the extent to which the association between cognition and ADHD symptoms was captured by the stacked model, we calculated the proportion of symptom-related variance in observed g-scores that was shared with the predicted g-scores. Specifically, this proportion was defined as the ratio of (a) the common variance in observed g-scores attributable to both a given ADHD symptom measure and the predicted g-scores to (b) the total variance in observed g-scores explained by that symptom measure.

We conducted these LME analyses in R 4.5.1, using lme4 (1.1-37) ^77^ for modelling fitting, lmerTest (3.1-3) ^78^ for model comparison with Satterthwaite degrees of freedom correction for fixed effects significance. Model comparisons used likelihood ratio tests via the anova() function, and confidence intervals were computed using the Wald method (broom.mixed, 0.2.9.6 ^79^). Variance decomposition used r2mlm (0.3.8)^37^, and commonality analyses used glmm.hp (0.1-8) ^42^. For figure presentation, components contributing less than 0.15% of the total variance were omitted for readability, as we determined they were not interpretable in this context. Full commonality tables including these components are reported in Supplementary Tables S8, S9, and S10.

### Confound sensitivity analyses

To assess the robustness of the principal findings to potential confounding factors, we conducted a series of sensitivity analyses examining the influence of sex, fMRI signal quality, sMRI signal quality, stimulant medication use, and sociodemographic status. Full details of the procedures and the rationale for confound selection are provided in the Supplementary Materials (Table S11).

Briefly, to evaluate potential sex differences, we tested both whether the predictive performance of the stacked model differed between males and females and whether it was statistically equivalent, using a test for the difference between two independent correlations together (Conventional difference Fisher’s Z test) with two one-sided tests (TOST) for equivalence, applied to Fisher z-transformed Pearson correlation ^80^

For the remaining potential confounds, we performed a residualisation analysis in which each confound, as well as all confounds combined, was regressed out of the stacked model’s predicted g-scores. We then re-evaluated the performance of the predicted g-scores in capturing both inter-individual and intra-individual variation in cognition, as well as their contributions to the variance decomposition and commonality analyses involving age and ADHD symptoms.

We operationalised fMRI signal quality as the number of volumes remaining after motion censoring, sMRI signal quality as the mean Euler number derived from FreeSurfer surface reconstruction, stimulant medication use as parent-reported medication status, and sociodemographic status as parental educational attainment.

## Results

### Confirmatory factor analysis (CFA) of cognitive functioning

A general cognitive factor (g) was derived from six cognitive measures using a first-order confirmatory factor analysis (CFA). Model fit indices averaged across cross-validation folds were as follows: comparative fit index (CFI; *M* = .929, range = .923–.942), Tucker– Lewis index (TLI; *M* = .882, range = .871–.903), root mean square error of approximation (RMSEA; *M* = .114, range = .103–.119), and standardized root mean square residual (SRMR; *M* = .043, range = .043–.049). Although several fit indices improved under a second-order CFA specification, the resulting g scores were nearly identical to those derived from the first-order model (*r* = .996, *p* < .0001), indicating that the operationalisation of g was highly robust to model specification (see Supplementary Materials).

The cognitive measures showed moderate-to-strong loadings on the latent g factor, with mean factor loadings ranging from .586 to .741. These loadings were highly consistent across cross-validation folds, supporting the stability and reproducibility of the derived g factor (Figure 1; Supplementary Table S4).

### Phenotype group differences in *g*

We next examined baseline differences in general cognitive ability (*g*) across phenotype groups at the first neuroimaging-aligned assessment. Mean *g* scores were −0.088 (*SD* = 0.80) for healthy controls, −0.633 (*SD* = 0.85) for participants with ADHD, and −0.350 (*SD* = 0.77) for the neither-ADHD-nor-clean-control group. Assumptions of normality and homogeneity of variance were met (Levene’s test: *F*(3, 590) = 1.34, *p* = .260). A one-way ANOVA revealed a significant effect of phenotype group on *g, F*(3, 590) = 20.04, *p* < .001, η^2^ = .092, ω^2^ = .088. Post hoc Tukey HSD tests showed that healthy controls had significantly higher *g* scores than participants with ADHD, Hedges’ *g* = 0.65, bootstrapped 95% CI [0.49, 0.83], *p* < .001.

### Machine-learning performance: First-stage models

We first trained machine learning models to predict *g* scores using each of seven sets of neuroimaging features and three algorithms (see Figure 2). For sets of neuroimaging features, Functional Connectivity had the highest out-of-sample Pearson’s *r* (*M*=.39), followed by ReHo (*M*=.32), ALFF (*M*=.29), Total Brain Volume (*M*=.24), Cortical Thickness (*M*=.20), Cortical Area (*M*=.19) and finally Subcortical Volume (*M*=.16), respectively. As for algorithms, Partial Least Squares regression (PLS) had the highest out-of-sample Pearson’s *r* (*M*=.27), followed by XGBoost (*M*=.25), and Kernel Ridge Regression (KRR) (*M*=.24).

**Fig. 2:**
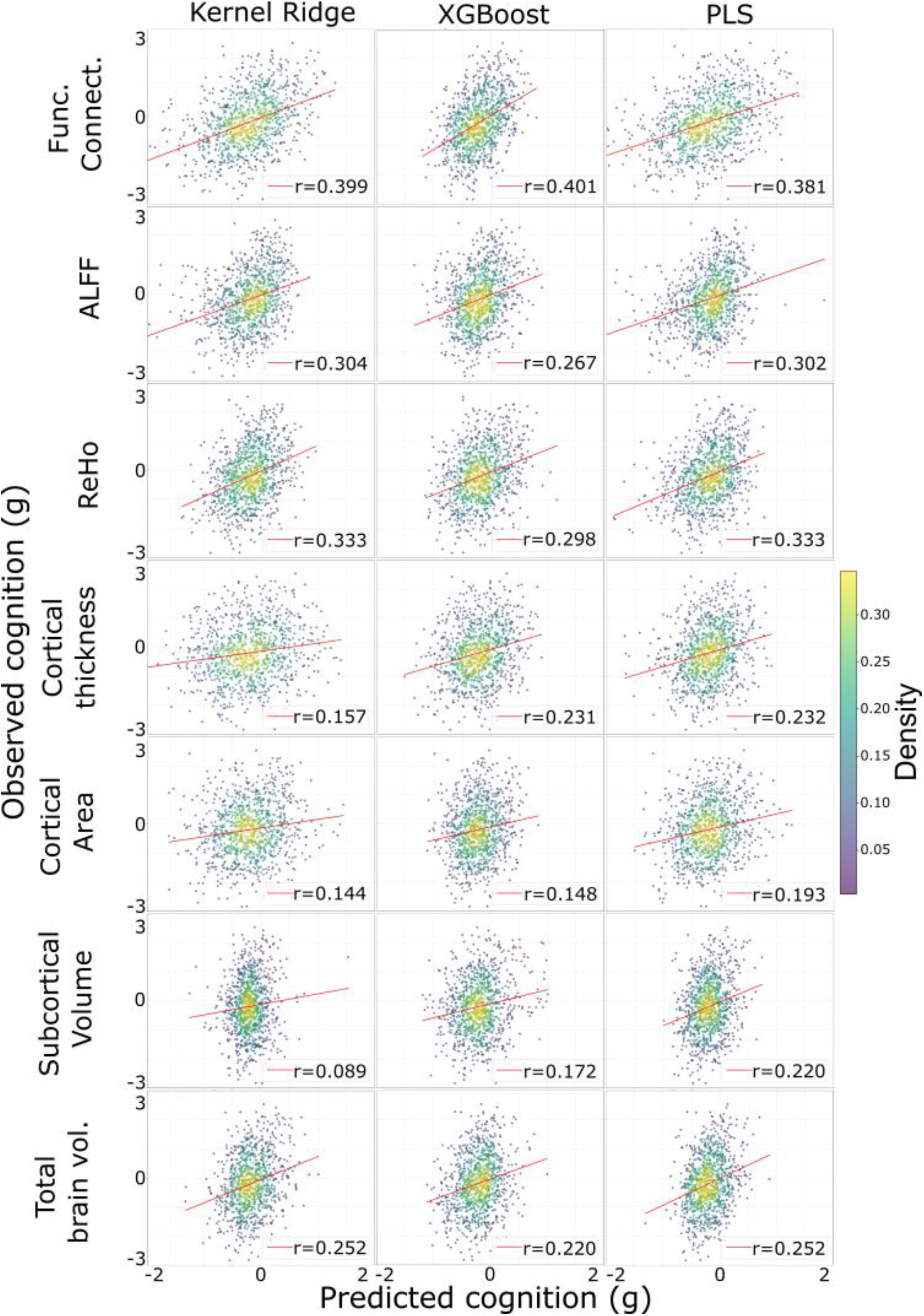
Scatterplots for observed vs. predicted g for each set of neuroimaging features and algorithms.

### Aim#1: Comparing stacked vs. first-stage models

The stacked model, which integrated information across multiple neuroimaging feature sets and algorithms to predict *g* scores, achieved an out-of-sample Pearson *r* of *M =* .459, an incremental but statistically significant improvement over the best-performing first-stage model (functional connectivity with XGBoost, *r* = .401 [0.025, 0.117]). Figure 3 shows a scatterplot of the stacked model performance, and Figure 4 shows bootstrapping analyses comparing the stacked model with each of the first-stage models.

**Fig. 3:**
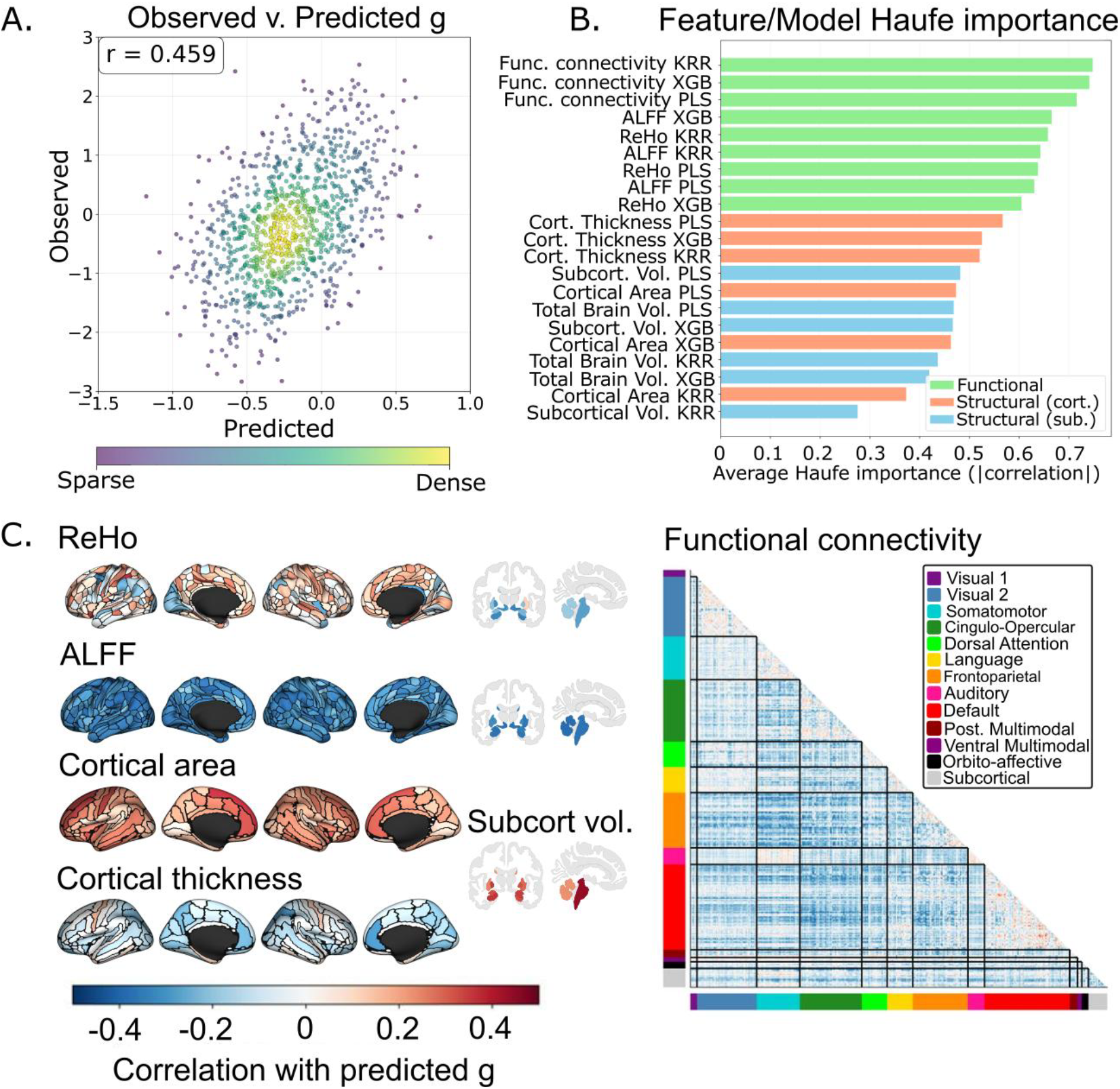
Feature importance and stacked model predictive performance. A) Scatter plot showing the correlation between observed and predicted g-scores, from the final stacked model (r= 0.459). B) Feature importance rankings (Haufe importance) across neuroimaging features and machine learning models, coloured by modality (green = functional, orange = structural - cortical only, and blue = structural subcortical or both cortical and subcortical). C) Brain surface/subcortical maps and functional connectivity matrix displaying the Pearson r between predicted g and regional brain features. Connectivity matrix (bottom right) is organised according to the functional network based on Cole-Anticevic organisation^81^, and represents the Pearson r between predicted g and Fisher’s Z connectivity.

**Fig. 4:**
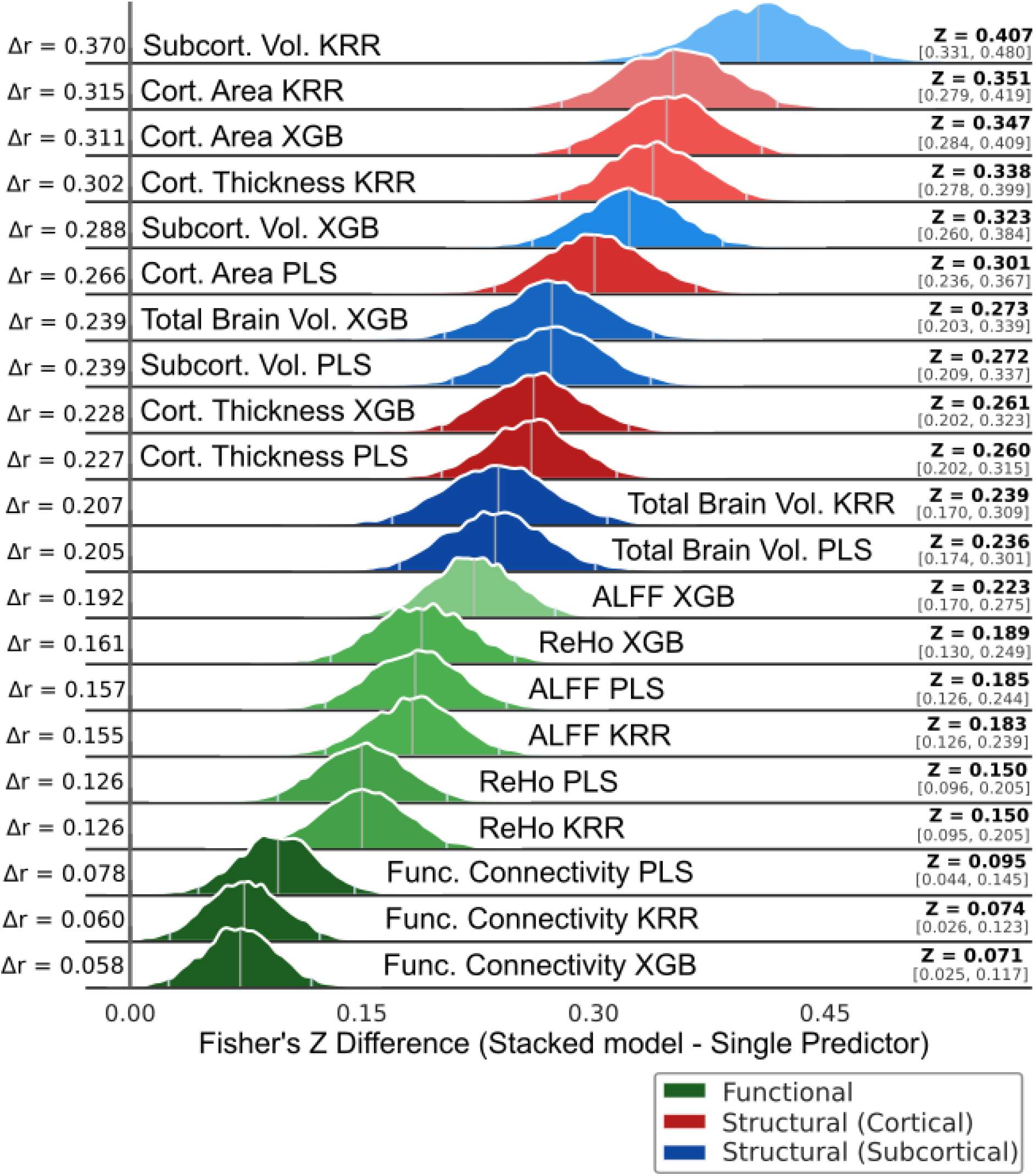
Bootstrap analysis comparing the stacked model with first-stage models. The x-axis represents the difference in Fisher Z-transformed correlations between the stacked model and each first-stage model (z-stacked minus z-first stage), such that the 95% confidence intervals higher than 0.00 indicate significantly better stacked model performance relative to first-stage model. The 0.00 line represents the null hypothesis of no difference between the stacked model and each first-stage model. Bootstrap distributions are hue coded by proximity to the 0.00 line (where greater distance is coded with lighter hue).

Based on feature-importance estimates (Figure 3B), fMRI-based first-stage models generally showed higher Haufe importance than sMRI-based first-stage models in terms of the association with stacked model predictions. Among all features, functional connectivity paired with Kernel Ridge Regression showed the highest mean Haufe importance (|*r* |=.763). For sMRI measures, cortical thickness was most closely related to stacked-model predictions, with Partial Least Squares yielding the highest mean Haufe importance (|*r* |=0.566).

Within the functional connectivity features, several large-scale networks showed relatively stronger associations with the stacked predictions, as indexed by the mean correlation across all region pairs within each network. The Frontoparietal Network exhibited the strongest association (*M* = −.14), followed by the Cingulo-Opercular Network (*M* = −.13) and the Default Mode Network (*M* = −.12) (Figure 3C, right panel). Full methodological details and the complete set of network-level correlations are provided in the Supplementary Materials and Table S7.

### ALFF and ReHo feature importance

Comparing our full stacked model with an ablated stacked model trained without ALFF and ReHo predictions, we found that the full stacked model (*r*=.459) outperformed the ablated stacked model (*r*=.429), with a mean Fisher’s Z difference of 0.038 (± 0.012 SD) in a bootstrapping analysis (95% CI: [0.014, 0.062]). This suggests that ALFF and ReHo led to a statistically significant, albeit small, improvement in predictive performance beyond Functional Connectivity and sMRI.

### Linear mixed-effects (LME) models

#### Generalisability of the predicted g-scores across ADHD diagnoses

To assess whether the stacked model predicted the g scores similarly for children with and without ADHD, we compared the ADHD-additive and ADHD-interactive models (see Methods). Both LME models regressed observed g-scores on predicted g-scores and ADHD status, with the ADHD-interactive model additionally including the predicted g-score × ADHD interaction term. As shown in Table 1A, the two models had comparable AIC and marginal R^2^ values, and the likelihood ratio test indicated that the ADHD-interactive model did not provide a better fit than the ADHD-additive model (X^2^(2, *N*=733)=0.73, *p*=.695). Because adding the interaction term did not improve model fit, the stacked model appears to predict the g-factor similarly for children with and without ADHD. Of note, however, the significant main effect of ADHD diagnosis in the additive model (β = -0.57, p<.001) indicates that children with ADHD scored lower on cognitive testing than controls, after accounting for neuroimaging-predicted *g*, which likely reflects aspects of ADHD-related cognition not fully captured by the s/fMRI marker.

**Table 1:** Linear Mixed Effects for ADHD diagnostic comparison, and Neuroimaging, Age, and Neuroimaging-and-Age models.

**A. ADHD Diagnosis Comparison**
| Predictors | ADHD Additive |  |  | ADHD Interactive |  |  |
| --- | --- | --- | --- | --- | --- | --- |
|  | Estimates | CI | p | Estimates | CI | p |
| (Intercept) | 0.44 | 0.28 – 0.59 | <0.001 | 0.44 | 0.29 – 0.60 | <0.001 |
| $\hat{g}_{\text{deviation}}$ | 0.37 | 0.32 – 0.43 | <0.001 | 0.37 | 0.29 – 0.46 | <0.001 |
| $\hat{g}_{\text{mean}}$ | 0.56 | 0.44 – 0.68 | <0.001 | 0.48 | 0.27 – 0.69 | <0.001 |
| ADHD Diagnosis | -0.57 | -0.77 – -0.38 | <0.001 | -0.58 | -0.77 – -0.38 | <0.001 |
| $\hat{g}_{\text{deviation}} \times$<br>ADHD | | | | -0.00 | -0.11 – 0.11 | 0.985 |
| $\hat{g}_{\text{mean}} \times$ ADHD | | | | 0.11 | -0.14 – 0.37 | 0.394 |
| **Random Effects** |  |  |  |  |  |  |
| $\sigma^2$ | 0.25 | | | 0.25 | | |
| $\tau_{00}$ | 0.53 | participant_id | | 0.53 | participant_id | |
| ICC | 0.68 |  |  | 0.68 |  |  |
| N | 279 | participant_id |  | 279 | participant_id |  |
| Observations | 733 |  |  | 733 |  |  |
| Marginal $R^2$ / Conditional $R^2$ | 0.297 / 0.775 | | | 0.298 / 0.775 | | |
| Deviance | 1578.920 |  |  | 1578.193 |  |  |
| AIC | 1607.011 |  |  | 1616.389 |  |  |

**B. Model Comparison: Neuroimaging and Age**
| Predictors | Neuroimaging Only |  |  | Age Only |  |  | Neuroimaging-and-Age |  |  |
| --- | --- | --- | --- | --- | --- | --- | --- | --- | --- |
|  | Estimates | CI | p | Estimates | CI | p | Estimates | CI | p |
| Intercept | 0.06 | -0.04 – 0.16 | 0.229 | -0.95 | -1.23 – -0.67 | <0.001 | -0.53 | -0.82 – -0.24 | <0.001 |
| $\hat{g}_{\text{deviation}}$ | 0.37 | 0.32 – 0.43 | <0.001 | | | | -0.01 | -0.09 – 0.07 | 0.782 |
| $\hat{g}_{\text{mean}}$ | 0.57 | 0.45 – 0.70 | <0.001 | | | | 0.44 | 0.31 – 0.58 | <0.001 |
| age<sub>deviation</sub> |  |  |  | 0.02 | 0.02 – 0.02 | <0.001 | 0.02 | 0.02 – 0.02 | <0.001 |
| age<sub>mean</sub> |  |  |  | 0.02 | 0.02 – 0.03 | <0.001 | 0.01 | 0.01 – 0.02 | <0.001 |
| **Random Effects** |  |  |  |  |  |  |  |  |  |
| $\sigma^2$ | 0.25 | | | 0.19 | | | 0.19 | | |
| $\tau_{00}$ | 0.60 | participant_id | | 0.69 | participant_id | | 0.59 | participant_id | |
| ICC | 0.71 |  |  | 0.78 |  |  | 0.76 |  |  |
| N | 283 | participant_id |  | 283 | participant_id |  | 283 | participant_id |  |
| Observations | 742 |  |  | 742 |  |  | 742 |  |  |
| Marginal $R^2$ / Conditional $R^2$ | 0.233 / 0.776 | | | 0.215 / 0.831 | | | 0.302 / 0.831 | | |
| Deviance | 1629.381 |  |  | 1525.169 |  |  | 1485.382 |  |  |
| AIC | 1652.516 |  |  | 1561.144 |  |  | 1533.743 |  |  |

ICC = Intraclass Correlation; AIC = Akaike Information Criterion. A) Linear mixed-effects models predicting observed g-scores from $\hat{g}_{\text{deviation}}$ , $\hat{g}_{\text{mean}}$ , and ADHD diagnosis (Additive model), with interaction terms between $\hat{g}_{\text{deviation}}/\hat{g}_{\text{mean}}$ and ADHD diagnosis additionally included in the Interactive model. Non-significant interaction
terms and comparable model fit indicate that the neuroimaging marker predicts $g$ similarly in children with and without ADHD; however, the significant main effect of ADHD diagnosis indicates a residual difference in observed $g$ not fully captured by the neuroimaging marker. B) Linear mixed-effects models predicting observed $g$ -scores from neuroimaging predictions alone ( $\hat{g}$ \_deviation, $\hat{g}$ \_mean; Neuroimaging Only), age alone (age\_deviation, age\_mean; Age Only), and both together (Neuroimaging-and-Age), used for commonality analysis (see Methods). The Neuroimaging-and-Age model showed improved fit over either single-predictor model (AIC = 1533.7 vs 1652.5 [neuroimaging] and 1561.1 [age]). The non-significant $\hat{g}$ \_deviation in this model reflects overlap between the intraindividual neuroimaging signal and age-related variance.

#### Aim#2: Decomposition of intra-versus inter-individual variation

To address our second aim, we evaluated the extent to which the stacked model captured variance in cognitive functioning at two levels: differences between individuals (inter-individual variation) and changes within individuals over time (intra-individual variation). To quantify the proportion of inter- and intra-individual variance in observed g scores explained by the stacked model, we conducted a variance decomposition analysis^75^ on the neuroimaging-only model (see Methods). Specifically, observed g scores were regressed on predicted g scores generated by the stacked model. To disentangle inter-individual and intra-individual components of the predictions, each participant’s mean predicted g score across all timepoints (ĝ_mean_, reflecting inter-individual variation) was calculated, along with the deviation of each timepoint-specific predicted score from that participant-specific mean (ĝ_deviation_ reflecting intra-individual variation).

Table 1B reports the beta estimates from this neuroimaging-only model, showing that both ĝ_mean_ and ĝ_deviation_ were significantly associated with observed g-scores. Figure 5 presents the variance decomposition, partitioning total variance in observed g-scores into interindividual and intraindividual components. The fixed effect of ĝ_mean_ accounted for 25.01% of the interindividual variance in observed g-scores (i.e., stable cognitive differences; denoted 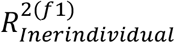, Figure 5E), consistent with the correlation between means of predicted and observed g-scores (*r*=.47, *p*<.0001, Figure 5C). Likewise, the fixed effect of ĝ_deviation_ accounted for 18.82 % of the intraindividual variance in observed g-scores (i.e., cognitive change within participants over time; denoted 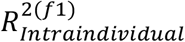, Figure 5E), consistent with the correlation between deviations of predicted and observed g-scores (*r*=.52, *p*<.0001, Figure 5D).

**Fig. 5:**
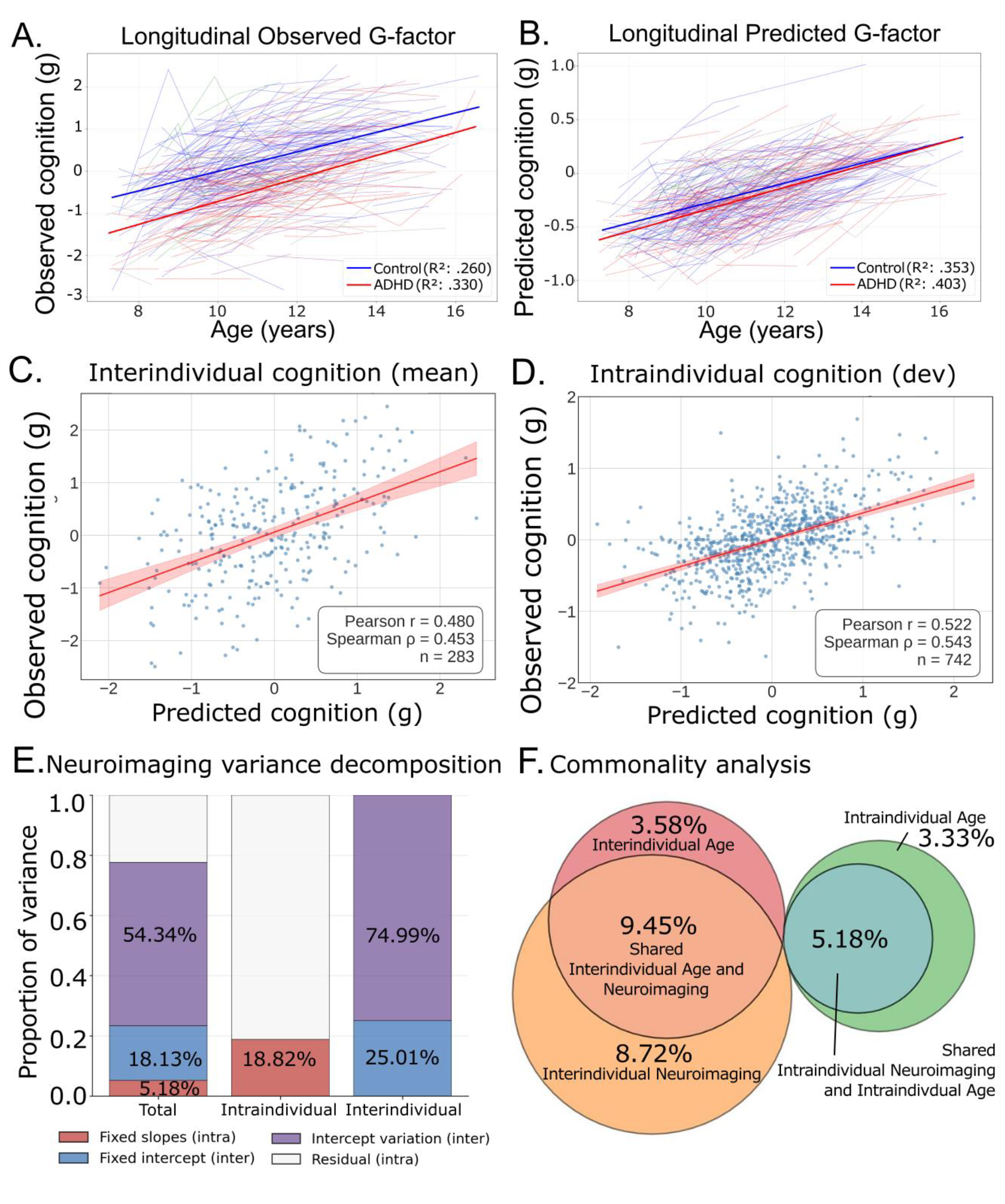
Longitudinal trajectories of observed and predicted cognition. A. Line plot showing the relationship between age and observed g-scores (derived from cognitive taasks) as a function of ADHD status. B. Line plot showing the relationship between age and predicted g-scores (derived from the stacked neuroimaging model) as a function of ADHD status. C. Scatter plot showing the relationship between mean predicted and mean observed g-scores, reflecting capacity of the stacked model in predicting inter-individual differences in cognition. D. Scatter plot showing the relationship between deviations in predicted and observed g-scores, reflecting capacity of the stacked model in predicting intraindividual differences in cognition. Dev depicts each participant’s deviation from their mean. E. Variance decomposition of the neuroimaging-only model. 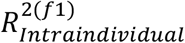, the proportion of intraindividual variance in observed g-scores explained by ĝ_deviation_, is shown in red in the middle panel. 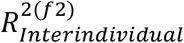, the proportion of interindividual variance explained by ĝ_mean_,, is shown in blue in the right panel. F. Euler diagram depicting the commonality analysis for the neuroimaging-and-age model. Percentages indicate the proportion of variance in observed g-scores explained jointly and uniquely by predicted g-scores and age components.

#### Aim#3: Commonality between age and predicted g-scores

Building on the finding that the neuroimaging marker captured both interindividual and intraindividual variance in cognitive functioning, we next asked how much of this captured variance reflected age-related cognitive development through commonality analyses^42,43^. Here, we used two additional LME models: an age-only model and a neuroimaging-and-age model (see Methods). The age-only model served as the baseline, regressing observed g-scores on age. As with ĝ_mean_ and ĝ_deviation_, we decomposed age into its interindividual component (age_mean_) and intraindividual component (age_deviation_). Table 1B presents the beta estimates from this age-only model, showing that both age_mean_ and age_deviation_ had a significantly positive association with observed g-scores. The neuroimaging-and-age model showed improved fit over the age-only model (AIC = 1533.7 vs 1561.1, ΔAIC = 27.4), indicating that the addition of the neuroimaging marker explained variance in cognitive functioning beyond that accounted for by age alone (Table 1B).

For the commonality analysis itself, we used the neuroimaging-and-age model, which regressed observed g-scores on ĝ_mean_, ĝ_deviation_, age_mean_, and age_deviation_. Table 1B shows that, when all four fixed-effect explanatory variables were included, the effect of ĝ_deviation_ was no longer significant, indicating overlap in the variance explained by ĝ_deviation_ and the age-related predictors.

The commonality analysis quantified this overlap (Figure 5F). For interindividual variability, the variance in observed g-scores attributable to age_mean_ consisted of 3.58% unique variance and 9.45% shared with ĝ_mean_, totaling 13.03%. This indicates that the stacked model captured (9.45/13.03)×100=72.52% of the interindividual age-related variance (see Supplementary materials for a detailed explanation of this calculation).

Similarly, for intraindividual variability, the variance in observed g-scores attributable to age_deviation_ consisted of 3.33% unique and 5.18% shared with ĝ_deviation_, totally 8.51%. This indicates that the stacked model captured (5.18/8.51)×100=60.87% of the intraindividual age-related variance. In other words, the stacked model tracked the majority of the cognitive development that would be expected from age alone, at both the interindividual and intraindividual level.

#### Aim#4: ADHD symptoms and cognition relationship

Having established that the neuroimaging marker tracked both cognitive differences between individuals and cognitive change within individuals, we next examined whether it also captured the relationship between cognitive functioning and ADHD symptoms. To assess the relationship between each ADHD symptom and observed g-scores, we used the hyperactivity-only and inattention-only models (see Methods). These models regressed observed g-scores on each of ADHD symptoms, derived from ADHD Rating Scale IV^44^. As with ĝ and age, we decomposed each ADHD symptom into its interindividual component (Hyperactivity_mean_ and Inattention_mean_) and intraindividual component (Hyperactivity_deviation_ and Inattention_deviation_).

Table 2 and Figure 6 show results from these ADHD symptom models. For hyperactivity, higher Hyperactivity_mean_ and Hyperactivity_deviation_ were significantly related to worse observed g-scores. In contrast, for inattention, while a higher Inattention_mean_ was significantly associated with worse observed g-scores, the effect of Inattention_deviation_ was not statistically significant. This suggests that interindividual variation in both hyperactivity and inattention was related to cognitive functioning, while only intraindividual changes in hyperactivity, but not inattention, were related to cognitive functioning.

**Table 2:**
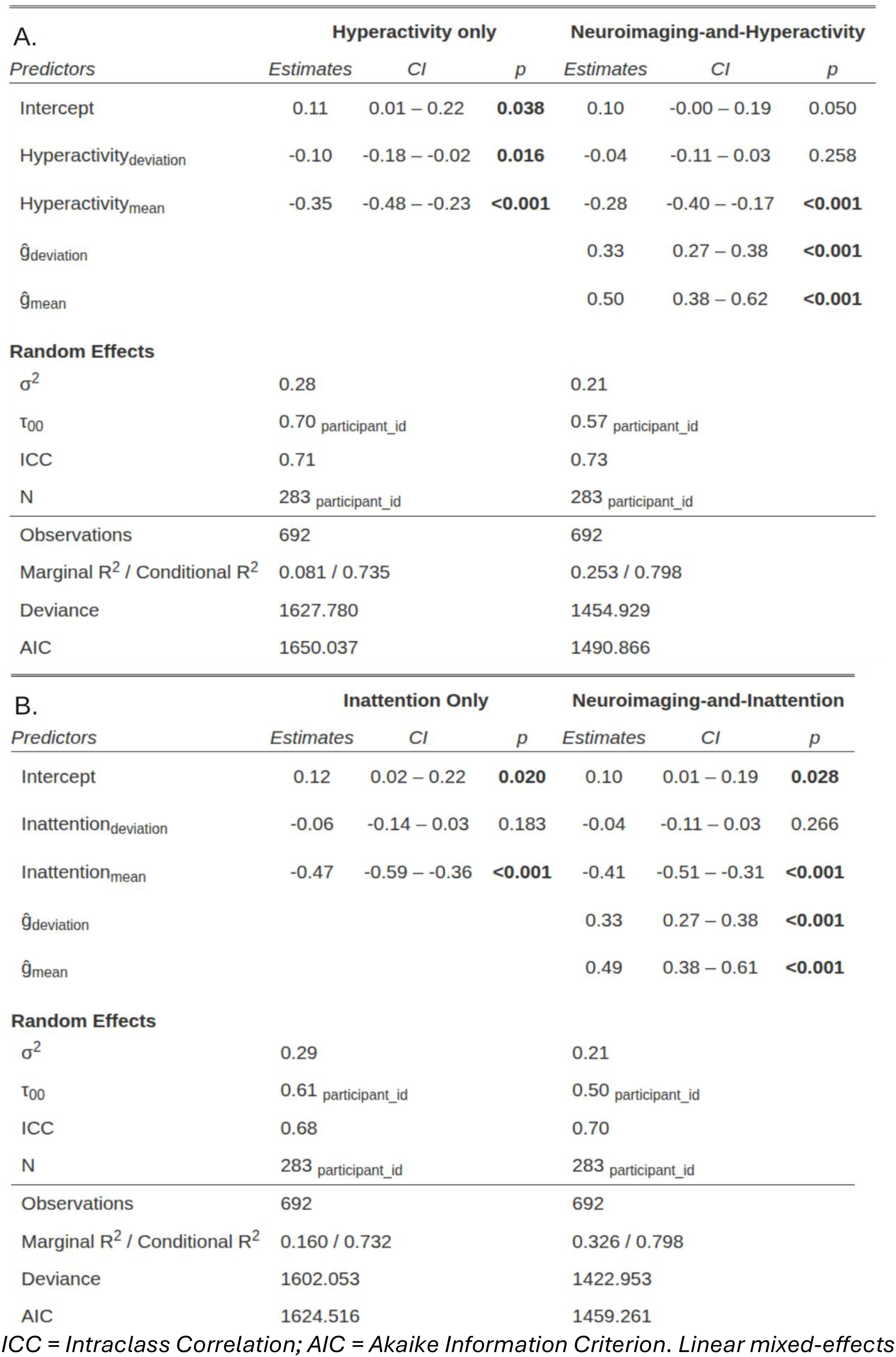
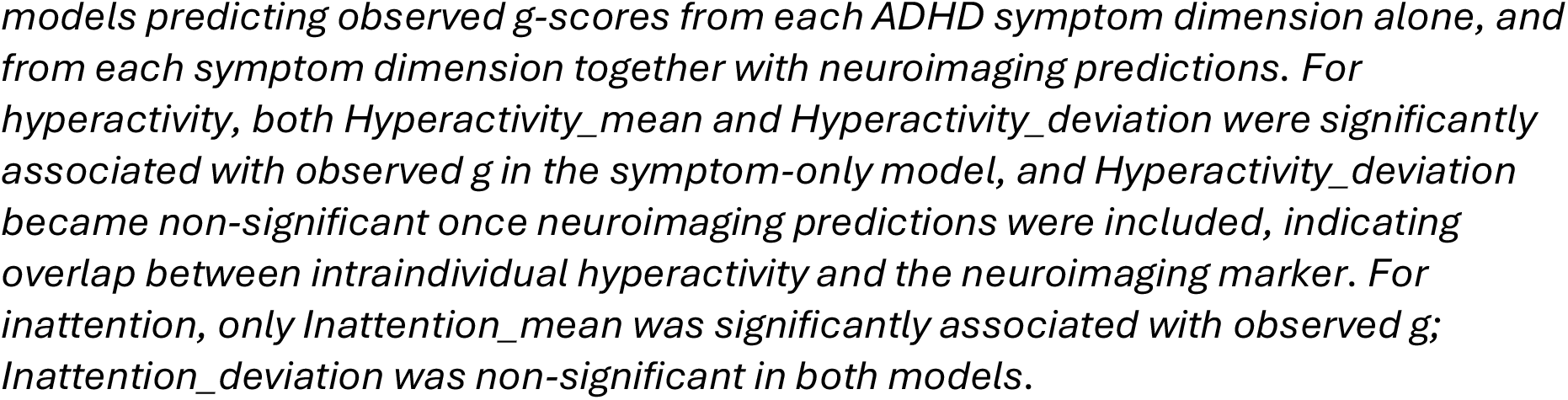
Linear mixed effects for Hyperactivity and Inattention symptoms, and neuroimaging for explaining variance in cognitive functioning.

| A. |  |  |  |  |  |  |
| --- | --- | --- | --- | --- | --- | --- |
| Predictors | Hyperactivity only |  |  | Neuroimaging-and-Hyperactivity |  |  |
|  | Estimates | CI | p | Estimates | CI | p |
| Intercept | 0.11 | 0.01 – 0.22 | **0.038** | 0.10 | -0.00 – 0.19 | 0.050 |
| Hyperactivity<sub>deviation</sub> | -0.10 | -0.18 – -0.02 | **0.016** | -0.04 | -0.11 – 0.03 | 0.258 |
| Hyperactivity<sub>mean</sub> | -0.35 | -0.48 – -0.23 | **<0.001** | -0.28 | -0.40 – -0.17 | **<0.001** |
| $\hat{g}_{deviation}$ | | | | 0.33 | 0.27 – 0.38 | **<0.001** |
| $\hat{g}_{mean}$ | | | | 0.50 | 0.38 – 0.62 | **<0.001** |
| **Random Effects** |  |  |  |  |  |  |
| $\sigma^2$ | 0.28 | | | 0.21 | | |
| $\tau_{00}$ | 0.70 | participant_id | | 0.57 | participant_id | |
| ICC | 0.71 |  |  | 0.73 |  |  |
| N | 283 | participant_id |  | 283 | participant_id |  |
| Observations | 692 |  |  | 692 |  |  |
| Marginal R<sup>2</sup> / Conditional R<sup>2</sup> | 0.081 / 0.735 |  |  | 0.253 / 0.798 |  |  |
| Deviance | 1627.780 |  |  | 1454.929 |  |  |
| AIC | 1650.037 |  |  | 1490.866 |  |  |
| B. |  |  |  |  |  |  |
| Predictors | Inattention Only |  |  | Neuroimaging-and-Inattention |  |  |
|  | Estimates | CI | p | Estimates | CI | p |
| Intercept | 0.12 | 0.02 – 0.22 | **0.020** | 0.10 | 0.01 – 0.19 | **0.028** |
| Inattention<sub>deviation</sub> | -0.06 | -0.14 – 0.03 | 0.183 | -0.04 | -0.11 – 0.03 | 0.266 |
| Inattention<sub>mean</sub> | -0.47 | -0.59 – -0.36 | **<0.001** | -0.41 | -0.51 – -0.31 | **<0.001** |
| $\hat{g}_{deviation}$ | | | | 0.33 | 0.27 – 0.38 | **<0.001** |
| $\hat{g}_{mean}$ | | | | 0.49 | 0.38 – 0.61 | **<0.001** |
| **Random Effects** |  |  |  |  |  |  |
| $\sigma^2$ | 0.29 | | | 0.21 | | |
| $\tau_{00}$ | 0.61 | participant_id | | 0.50 | participant_id | |
| ICC | 0.68 |  |  | 0.70 |  |  |
| N | 283 | participant_id |  | 283 | participant_id |  |
| Observations | 692 |  |  | 692 |  |  |
| Marginal R<sup>2</sup> / Conditional R<sup>2</sup> | 0.160 / 0.732 |  |  | 0.326 / 0.798 |  |  |
| Deviance | 1602.053 |  |  | 1422.953 |  |  |
| AIC | 1624.516 |  |  | 1459.261 |  |  |

ICC = Intraclass Correlation; AIC = Akaike Information Criterion. Linear mixed-effects
models predicting observed *g*-scores from each ADHD symptom dimension alone, and from each symptom dimension together with neuroimaging predictions. For hyperactivity, both *Hyperactivity\_mean* and *Hyperactivity\_deviation* were significantly associated with observed *g* in the symptom-only model, and *Hyperactivity\_deviation* became non-significant once neuroimaging predictions were included, indicating overlap between intraindividual hyperactivity and the neuroimaging marker. For inattention, only *Inattention\_mean* was significantly associated with observed *g*; *Inattention\_deviation* was non-significant in both models.

**Fig 6.**
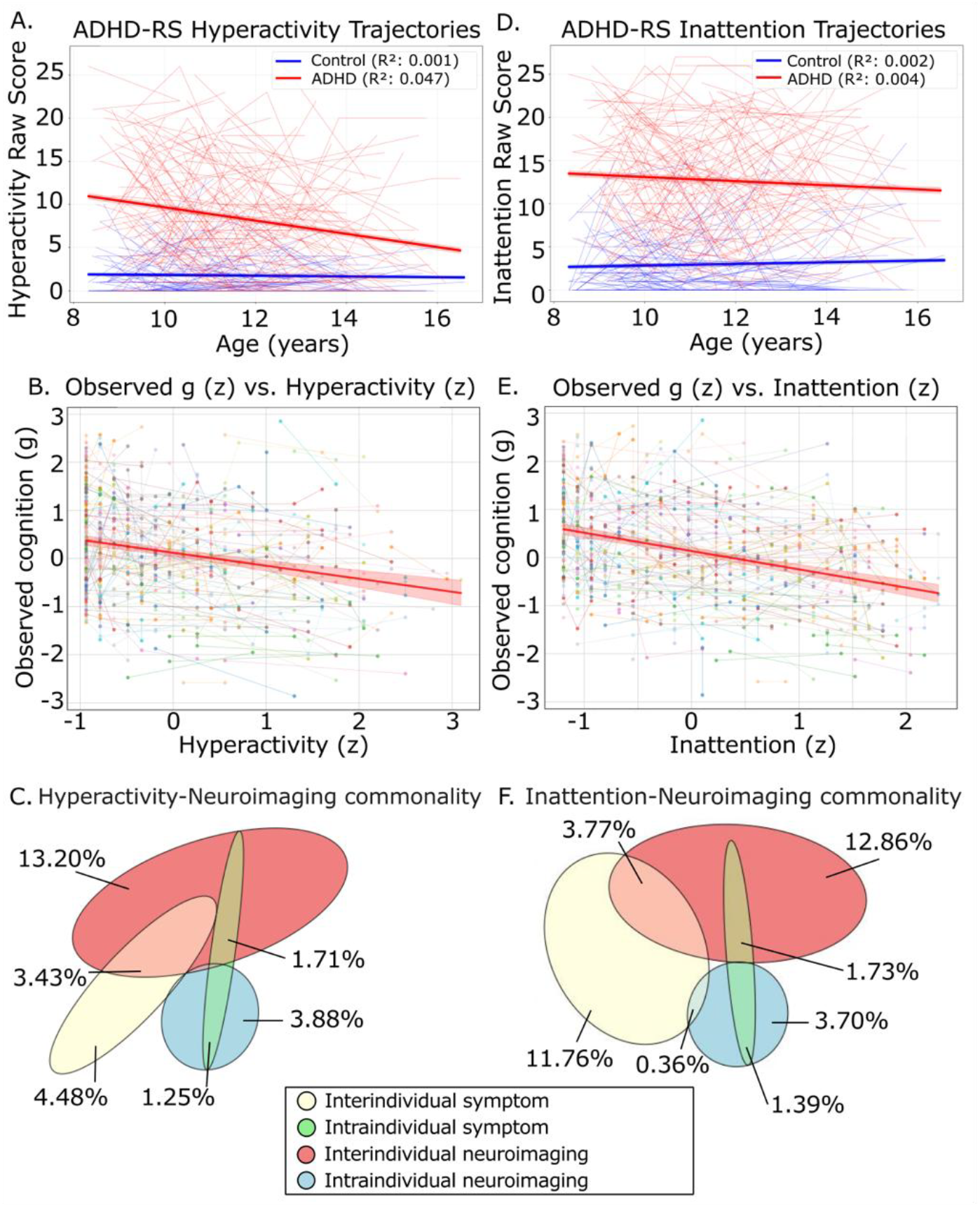
Symptom and cognition relationships and the commonality between ADHD symptoms and predicted g-scores. A. Hyperactivity trajectories of participants with more than one neuroimaging timepoint. B. Scatter plot showing the relationship between hyperactivity and observed g-score C. Euler diagram depicting the commonality analysis for the neuroimaging-and-hyperactivity model. Percentages indicate the proportion of variance in observed g-scores explained jointly and uniquely by predicted g-scores and hyperactivity. (note commonality between factors less than 0.15% not reported here, see supplementary) D. Inattention trajectories of participants with more than one neuroimaging timepoint. E. Scatter plot showing the relationship between inattention and observed g-score F. Euler diagram depicting the commonality analysis for the neuroimaging-and-inattention model. Percentages indicate the proportion of variance in observed g-scores explained jointly and uniquely by predicted g-scores and inattention. (note commonality between factors less than 0.15% not reported here, see supplementary)

### Commonality between ADHD symptoms and predicted g-scores

To evaluate how well the stacked model captured the relationship between cognition and ADHD symptoms, we conducted commonality analyses^42,43^ using the neuroimaging-and-hyperactivity and neuroimaging-and-inattention models (see Methods).

#### Neuroimaging-and-hyperactivity model

The neuroimaging-and-hyperactivity model regressed observed g-scores on ĝ_mean_, ĝ_deviation_, Hyperactivity_mean_, and Hyperactivity_deviation_. Table 2 shows that, when all four fixed-effect explanatory variables were included, Hyperactivity_deviation_ was no longer statistically significant, indicating that its explanatory variance overlapped with that of on ĝ_mean_ and/or ĝ_deviation_. The commonality analysis quantified this overlap (Figure 6).

The variance in observed g-scores attributable to Hyperactivity_mean_ consisted of 4.48% unique variance and 3.43% shared with ĝ_mean_, totalling 7.91%. This indicates that the stacked model captured (3.43/7.91)×100=43.36% of the interindividual variability in hyperactivity.

The variance in observed g-scores attributable to Hyperactivity_deviation_ comprised 1.71% shared with ĝ_mean_ and 1.25% shared with ĝ_deviation_, equating to 2.96%, with no unique component. This shows that the stacked model captured 100% of the intraindividual variability in hyperactivity. Notably, however, this capture was limited in absolute terms (2.96% of the shared variance in g).

Overall, the stacked model captured (3.43+1.25+1.71)/(4.48+3.43+1.25+1.71)×100=58.79% of the total variability in hyperactivity modelled by our variables (6.39% of total variance in g).

#### Neuroimaging-and-inattention model

The neuroimaging-and-inattention model regressed observed g-scores on ĝ_mean_, ĝ_deviation_, Inattention_mean_, and Inattention_deviation_. Table 2 shows that, when all four predictors were included, Inattention_deviation_ was not statistically significant. Unlike Hyperactivity_deviation_, however, Inattention_deviation_ was not statistically significant even when predicted g-scores were excluded. Therefore, our commonality analysis for inattention focused only on Inattention_mean_.

As seen in Figure 6, the variance in observed g-scores attributable to Inattention_mean_ comprised 11.76% unique variance, 3.77% shared with ĝ_mean,_ and .36% shared with ĝ_deviation_, totalling 15.89%. This indicates that the stacked model captured (3.77+.36)/15.89×100=25.99% of the interindividual variability in inattention (4.13% of total variance in g).

### Confound sensitivity analysis results

See the Supplementary Materials and Supplementary table 11 for the full results of the confound sensitivity analyses. Briefly, the predictive performance of the stacked model did not differ significantly between males (r = .438) and females (r = .491; z = -1.04, p = .30) and was statistically equivalent across sexes (p = .040), indicating comparable performance for male and female participants. Residualising fMRI signal quality (indexed by the number of volumes remaining after censoring) attenuated inter-individual prediction (r = .35 vs. .47 before residualisation) but had virtually no effect on intra-individual prediction (r = .52 both before and after residualisation). Furthermore, residualising sMRI signal quality (indexed by FreeSurfer’s mean Euler number), stimulant medication use, and sociodemographic status (indexed by parental education) did not reduce predictive performance for either inter-individual prediction (r = .49, .49, and .48, respectively, vs. .47 before residualisation) or intra-individual prediction (r = .53, .53, and .54, respectively, vs. .52 before residualisation). Collectively, these findings indicate that the ability of the stacked sMRI/fMRI marker to predict within-person changes in cognitive functioning is robust to all tested potential confounds.

## Discussion

Guided by the RDoC framework^15^, our findings addressed four primary aims. First, we evaluated whether multimodal sMRI and fMRI measures could serve as neurobiological markers of cognitive functioning in children with and without ADHD. Integrating diverse structural and functional features, including ReHo and ALFF, improved out-of-sample prediction and yielded a marker that generalized across diagnostic groups. Second, consistent with the developmental focus of RDoC^15^, we examined whether this marker tracked within-person cognitive changes over time. The marker explained 25.01% (*r* = .48) of between-person differences and 18.82% (r = .52) of within-person cognitive changes. Third, we assessed the extent to which neuroimaging features accounted for age-related variation in cognition. Commonality analyses showed that these features explained 72.52% of interindividual age differences and 60.87% of intraindividual age-related change, corresponding to 9.45% and 5.18% of the total variance in observed g-scores, respectively. Finally, given the relevance of RDoC markers to psychopathology^17^, we examined cognition–ADHD symptom associations. The marker explained 58.79% of the cognition–hyperactivity association and 25.99% of the cognition–inattention association, accounting for 6.39% and 4.13% of the total variance in observed g-scores, respectively.

### Aim#1: Combining sMRI and fMRI through multimodal stacking improved the prediction of cognitive functioning in ADHD

The RDoC framework conceptualises cognitive functioning as a core functional domain and emphasises its assessment across multiple neurobiological units of analysis^17^. Consistent with this framework, we tested whether a marker integrating diverse sMRI and fMRI features, including often-overlooked resting-state measures such as ReHo and ALFF, could capture cognition in individuals with and without ADHD. Multimodal integration improved prediction of cognitive functioning, consistent with previous work^31–33^. Functional connectivity was the set of features most relevant to prediction, reflecting the prominent role of large-scale brain networks in cognition^82^. The predictive strength of functional connectivity was more related to the frontoparietal network (mean r = -.14), aligning with the theoretical relevance of the frontoparietal network for cognitive functioning ^83^; the cingulo-opercular network (mean r = -0.13), where efficiency across childhood and adolescence is predictive of cognitive functioning in task performance ^84^; and the default mode network (r = -0.12), where inter-network connectivity is implicated in semantic and episodic memory, and attentional processes^85^.

Importantly, ReHo^26^ and ALFF^27^ provided significant incremental predictive value. Although their contributions were modest, these measures can be derived from standard resting-state fMRI data without additional acquisition costs, supporting their inclusion in future neuroimaging marker development. The resulting multimodal marker also generalised to children with ADHD. Its predictive performance (r = .459) in this ADHD-enriched cohort of young children is comparable to that reported in population-based studies^23–25^ and a meta-analysis of healthy adults (r = .42)^23^. Furthermore, formal statistical tests using ADHD interaction models revealed no significant moderation by ADHD status, indicating that the relationship between the neuroimaging marker and cognitive functioning was similar across children with and without ADHD. This finding is consistent with recent evidence from the population-based Adolescent Brain Cognitive Development (ABCD) Study, which demonstrated the generalizability of structural and functional MRI markers to children with ADHD within the study^18,22^. Importantly, all reported performance estimates were derived from predictions on held-out test data within a nested cross-validation framework, making it unlikely that the observed predictive accuracy reflects overfitting. If substantial overfitting were present, we would expect to observe markedly higher performance in the training data than in the held-out test data, with consistently poorer generalization to the test sets. This pattern was not observed in our analyses.

### Aim#2: The multimodal neuroimaging markers explained intra-individual (within-person) variation in cognitive functioning

The RDoC framework emphasises that functional domains are inherently developmental and should be assessed across time^15^. Accordingly, for a multimodal neuroimaging marker to serve as an RDoC-informed marker of cognitive functioning, it should be sensitive not only to between-person differences but also to within-person developmental change^32,34^. Although resting-state fMRI has historically been criticised for low test–retest reliability ^35^, our findings add to growing evidence ^32,33,36^ that machine-learning approaches integrating structural and functional neuroimaging features can yield reliable and developmentally informative markers.

The ability to capture longitudinal changes is particularly noteworthy because developmental prediction is rarely evaluated in machine-learning studies of ADHD, which remain predominantly cross-sectional^86^. While explaining 18.82% of within-person cognitive trajectories may appear modest in absolute terms, the corresponding intra-individual correlation (*r* = .52) represents a large effect in developmental research, exceeding the effect sizes reported in the majority of developmental psychology studies and falling at approximately the 75th percentile or higher of the empirical effect-size distribution ^87^. These findings suggest that multimodal neuroimaging markers can capture meaningful developmental variation in cognitive functioning, supporting their utility for longitudinal investigations of neurocognitive development in ADHD and typical populations.

### Aim#3: The multimodal neuroimaging markers accounted for age-related variation in cognition

To demonstrate that s/fMRI-based markers capture the developmental nature of RDoC functional domains, rather than merely tracking nonspecific fluctuations in brain state, it is necessary to show that they account for age-related cognitive development. Age is a particularly important reference point in ADHD research because cognitive performance generally improves across adolescence in both youth with ADHD and their typically developing peers^38–41^. Our multimodal marker accounted for 60.87% (5.18% of total variance in g) of within-person, age-related cognitive changes. We found that our multimodal marker accounted for 60.87% of within-person age-related cognitive change, corresponding to 5.18% of the total variance in g. Notably, the marker’s intraindividual contribution to cognitive functioning was entirely shared with age, such that it no longer explained unique variance in g after age was included in the model.

This finding is informative because the machine-learning models were trained exclusively on neuroimaging features and were never provided age information. The overlap therefore suggests that the within-person cognitive signal derived from brain features tracks the same developmental changes indexed by age. In a sample spanning ages 8–17 years, a period characterised by robust and well-documented structural and functional brain maturation^88–91^, the most parsimonious interpretation is that the marker’s sensitivity to cognitive change is driven by underlying neurodevelopmental processes. Although future studies are needed to directly test this interpretation, the present findings indicate that the marker captures cognitive variation closely aligned with normative developmental change. Indeed, within this age range, age may be best conceptualised as a proxy for ongoing neurodevelopment, providing a meaningful benchmark against which to evaluate the developmental relevance of neuroimaging-based cognitive markers.

This inference supports the use of s/fMRI for characterising developmental trajectories in relation to cognitive maturation in ADHD, but highlights the need to disentangle broadly normative neurodevelopment from any potential ADHD specific deviation from it as an important future direction. Our study utilised a supervised model to predict overall cognitive functioning; here, the brain signal and age are collinear, and there is no reference frame against which to define an individual’s departure from the expected developmental course. Further, lines of best fit through ADHD and control groups (see Figure 5) may be broadly similar while collapsing precisely the individual variation in cognitive trajectory that may constitute cognitive heterogeneity in ADHD. Recovering that variation would require a different lens. Future research could navigate this challenge through normative modelling frameworks^92,93^, which quantify individual deviations from reference population norms and may capture any ADHD-specific trajectories more sensitively than predicting absolute cognitive values.

### Aim#4: The multimodal neuroimaging markers accounted for ADHD-symptom-related variation in cognition

To establish the clinical relevance of the s/fMRI-based cognitive marker, and thereby link cognitive functioning as an RDoC domain to neuroimaging as its unit of analysis, we examined the extent to which the marker accounted for associations between cognitive functioning and ADHD symptoms. Consistent with previous research^45–48^, cognitive functioning was significantly associated with both hyperactivity and inattention. However, our longitudinal analyses revealed important distinctions between symptom domains. Hyperactivity was associated with cognitive functioning at both the between-person and within-person levels, indicating that individuals with greater hyperactivity tended to show lower cognitive functioning and that changes in hyperactivity over time tracked changes in cognition. In contrast, inattention was associated with cognitive functioning only at the between-person level. The absence of a within-person association is consistent with the relative stability of inattention across the age range examined in the present study (Fig. 6D), which limits the extent to which intraindividual fluctuations in attention symptoms can covary with cognitive change.

We then showed that our multimodal neuroimaging marker successfully captured these cognitive-and-symptom links. For the cognitive–hyperactivity association, our marker accounted for a substantial portion both inter-individually (43.36%; 3.43% of total variance in *g*) and intra-individually (100%). Although the intra-individual association between cognitive functioning and hyperactivity was modest in absolute terms (accounting for 2.96% of the variance in *g*), our marker captured its full extent. For the cognitive–inattention relationship, the marker accounted for 25.99% of the inter-individual association (4.13% of total variance in *g*). This capacity positions our marker as a promising candidate for an RDoC-aligned cognitive-system biomarker^17^ that captures cognition–psychopathology links across development, spanning from typical to highly atypical.

### Confound sensitivity analyses

We conducted sensitivity analyses to evaluate the potential influence of five confounds: sex, fMRI signal quality, sMRI signal quality, stimulant medication use, and sociodemographic status (see Supplementary Materials). Of these, only fMRI signal quality affected predictive performance. Specifically, residualising the number of fMRI volumes remaining after motion censoring, a proxy for fMRI data quality, reduced between-person prediction accuracy (*r* = .35, compared with *r* = .47 before residualisation). This finding is consistent with evidence that predictive performance is influenced by the amount of usable fMRI data ^94^, highlighting the importance of acquiring sufficiently long scans in paediatric samples to ensure adequate data following motion censoring.

Importantly, adjustment for fMRI signal quality did not meaningfully affect within-person prediction (*r* = .52 both before and after residualisation). Thus, the marker’s ability to track longitudinal changes in cognitive functioning remained intact after accounting for data quality. Moreover, none of the five confounds substantially altered the performance of the stacked s/fMRI marker, indicating that its capacity to predict within-person cognitive change was robust to variation in sex, imaging quality, stimulant medication use, and sociodemographic factors.

## Limitations

Several limitations warrant acknowledgement. First, the sample averaged only 1.77 time points per participant, with many participants contributing just two observations. Consequently, intraindividual estimates were driven largely by two-point within-person contrasts, spaced on average 23 months apart, rather than by dense and evenly spaced developmental trajectories. This sparse longitudinal structure limited our ability to estimate random slopes reliably: linear mixed-effects models including random slopes failed to converge, likely because too few within-person observations were available to distinguish participant-specific slopes from residual variation^77,95^. Given that commonality analyses required us to maintain the same random-effects structure across models to facilitate model comparisons, we followed the keep it maximal principle^74^, and restricted the random-effects structure to random intercepts only. As a result, estimates of intraindividual change were derived from a relatively parsimonious longitudinal framework and should be interpreted accordingly.

Next, while we used relatively large sample of children and adolescents with ADHD in the current study, the present study did not evaluate the extent to which the s/fMRI markers identified here generalise across independent datasets. Future research should directly assess the external validity of these neuromarkers by examining their reproducibility across the Oregon ADHD-1000, longitudinal subsets of the ABCD Study ^96,97^, and other longitudinal ADHD cohorts such as NeuroIMAGE^98^. Such work would provide an important test of the robustness and generalisability of the identified s/fMRI markers.

Another important limitation is related to stimulant medication. The medication was discontinued 48 hours prior to assessment in the Oregon ADHD 1000 study, reducing the likelihood that our findings reflect acute medication effects. Nevertheless, potential longer-term influences of sustained stimulant treatment on brain–cognition relationships cannot be entirely ruled out^99^. Importantly, our findings remained robust after residualising medication effects at both the interindividual level (original *r* = .47; medication-residualised *r* = .49) and the intraindividual level (original *r* = .52; medication-residualised *r* = .54). Full results are provided in the Supplementary Materials (Table S11).

Lastly, we focused on developing s/fMRI markers of general cognitive ability (g), despite the RDoC framework distinguishing multiple cognitive domains, including attention, perception, declarative memory, language, cognitive control, and working memory^17^. We chose to focus on g for several reasons. First, our primary aim was prediction, and in the past we found that deriving a g factor from multiple cognitive tasks using factor analysis yielded substantially better predictive performance than modelling individual cognitive tasks separately ^100^. This likely reflects the higher reliability and signal-to-noise ratio of a latent factor relative to task-specific performance measures. Second, the cognitive battery available in the Oregon ADHD-1000 dataset does not comprehensively capture all RDoC cognitive constructs, particularly domains such as multisensory perception. Nevertheless, developing neuroimaging markers for specific cognitive domains beyond general cognitive ability represents an important direction for future RDoC-inspired research.

## Conclusion

Guided by the RDoC framework, we developed a multimodal MRI-derived marker of cognitive functioning in ADHD. The marker captured not only interindividual differences in cognition but also longitudinal cognitive changes, and accounted for meaningful associations between cognition and ADHD symptoms. Although predictive performance remains below the threshold required for clinical application, our findings demonstrate the feasibility of using structural and resting-state functional MRI to track cognitive functioning across development in ADHD. More broadly, these results provide a foundation for future RDoC-informed investigations of cognitive development and heterogeneity in ADHD. Future research could extend this work by incorporating additional neuroimaging modalities, such as task-based fMRI, which often yields stronger brain–cognition associations than resting-state measures^32,68^, as well as other RDoC-relevant units of analysis, including genomic, molecular, and physiological factors. Integrating these approaches with environmental measures may further advance understanding of the developmental processes that shape cognitive functioning in ADHD.

## Acknowledgements

Data and/or research tools used in the preparation of this manuscript were obtained from the National Institute of Mental Health (NIMH) Data Archive (NDA). NDA is a collaborative informatics system created by the National Institutes of Health to provide a national resource to support and accelerate research in mental health. Dataset identifier(s): The Oregon ADHD-1000: a longitudinal data resource enriched for clinical cases and multiple levels of analysis (Study ID#1938, DOI:10.15154/1528485). This manuscript reflects the views of the authors and may not reflect the opinions or views of the NIH, the Oregon ADHD-1000 researchers, or of those submitting original data to NDA.

K. Jack Scott was supported by a University of Otago Research Grant (0125-0326). Narun Pat was supported by Health Research Council of New Zealand (grant numbers 21/618 and 24/838), by Neurological Foundation of New Zealand (grant number 2350 PRG), and by the Ministry of Business, Innovation and Employment (grant numbers UOA2421 and RTVU2403). Kseniia Konopkina, Farzane Lal Khakpoor and Irina Buianova were all supported by a University of Otago Doctoral Scholarship. William van der Vliet was supported by Health Research Council of New Zealand (grant numbers 21/618 and 24/838).

## Code availability and data access

All code and scripts used in these analyses are publicly available at our laboratory github page: https://github.com/HAM-lab-Otago-University/Longitudinal-Multimodal-MRI-neuromarkers-ADHD/tree/main

The Oregon ADHD-1000 is available at the NDA NIMH Data archive: 10.15154/1528485

