## Supplementary Materials for "Multimodal Magnetic Resonance Imaging-based neuromarkers trace longitudinal changes in cognitive functioning in Attention Deficit/Hyperactivity Disorder"

### **Participant Race and Ethnicity data**

**Table S1.**
*Ethnicity composition of the final sample*

| Race | Non-Hispanic/Latino | Hispanic/Latino | Total |
| --- | --- | --- | --- |
| Asian | 9 | - | 9 |
| Black or African American | 8 | 1 | 9 |
| Hawaiian/Pacific Islander | - | - | 0 |
| Native American (or Alaskan) | 1 | - | 1 |
| White | 428 | 24 | 452 |
| More than one race | 54 | 6 | 60 |
| Not specified |  |  | 63 |

### **Cognitive battery and variable selection**

From the Oregon-1000 cognitive and psychometric testing battery, we selected the following cognitive tests: N-Back (a measure of working memory and attention)^1^, The Continuous Performance Task (CPT; a measure of sustained attention and vigilance)^2^, The Stop Signal Reaction Time Task (SSRT; measuring cognitive control and response inhibition)^3^, Digit Span Forward (DSF), and Digit Span Backward (DSB) task components of the Wechsler Intelligence Scale for Children IV^4^ (WISC-IV), measuring working memory for digit span forward, and working memory and cognitive control for digit span backward^5^, respectively. We also assessed two additional domains from the Wechsler Individual Achievement Test^6^ (WIAT): word reading score and maths reasoning score.

Cognitive tests were selected on the basis of data coverage aligned with our neuroimaging data (within 6 months of each scan time point). Data coverage is presented in Table S1 below.

We selected variables from these cognitive tests that were known to accurately capture the cognitive constructs of interest. We selected 2-back accuracy from the N-Back task, as a ceiling effect was observed with the majority of participants scoring near 100% accuracy in the 0-back condition, rendering a 2-back/0-back contrast effectively a 2-back accuracy.

**Table S2.**
*Data coverage for cognitive and symptom testing by phenotype group and test.*

| Task | Overall coverage | Control | Subthreshold | ADHD | Other | Not Assessed |
| --- | --- | --- | --- | --- | --- | --- |
| Stop Signal Reaction Time Task | 99.6% | 99.4% | 100% | 99.8% | 100% | 99.4% |
| Continuous Performance Task | 96.1% | 97.6% | 96.3% | 93.3% | 100% | 96.4% |
| N-Back | 96.6% | 97.6% | 95.7% | 95.2% | 100% | 95.8% |
| WISC-IV Digit span task | 87.6% | 85.8% | 87.6% | 88.1% | 75.4% | 97.6% |
| WIAT word reading score | 99.7% | 99.8% | 99.4% | 99.6% | 100% | 100% |
| ADHD-Rating Scale IV | 89.7% | - | - | - | - | - |

*Note: ADHD-RS-IV scores were used to determine phenotype classification, and therefore coverage reflects the broader clinical assessment schedule that aligns with neuroimaging timepoints.*

**Table S3.**
*Cognitive tests and measures utilised in deriving g-factor scores and correlation with other tests*

| Task | Cognitive domain | Measure(s) used | Mean Pearson r |
| --- | --- | --- | --- |
| Stop Signal Reaction Time Task | Cognitive control/response inhibition | Standard deviation of go signal reaction time | 0.4 |
| Continuous Performance task | Sustained Attention/Vigilance | D’Prime catch trials | 0.49 |
| N-Back | Working memory/Attention | 2-Back accuracy (%) | 0.43 |
| WISC-IV Digit span task | Working memory/cognitive control | DSF, DSB scores | DSF: 0.44, DSB: 0.43 |
| WIAT Word Reading | Reading comprehension/decoding | Word reading score | 0.4 |

*Mean correlation between each variable and all other variables across each participant-timepoint (Pearson’s r).*

**General cognitive factor (g-factor)**

To aid model convergence, for each fold, we calculated the sample covariance matrix and added a small regularisation constant (0.05) to the diagonal to improve numerical stability and thus model convergence^7,8^. We then fitted CFA models using full information maximum likelihood estimation (FIML) with standardised latent variables, and non-linear minimisation subject to box constraints for optimisation. We selected FIML to handle any potential missingness in data, and fixed the variance of latent to 1.

**Table S4**
*Standardised factor loading means for each cognitive variable, and range (across folds)*

| Variable | Mean factor loading | Range of factor loading |
| --- | --- | --- |
| CPT d’ prime catch | 0.741 | 0.715 – 0.788 |
| Digit Span forward | 0.644 | 0.632 – 0.663 |
| Digit Span backward | 0.635 | 0.619 – 0.650 |
| N-Back 2-back accuracy | 0.628 | 0.613 – 0.637 |
| SST reaction time SD | 0.591 | 0.570 – 0.611 |
| WIAT Word Reading score | 0.586 | 0.564 – 0.603 |

### **General cognitive factor CFA sensitivity analysis**

To evaluate whether the derived general cognitive factor (*g*) depended on the specific CFA specification, we conducted a sensitivity analysis comparing first-order and second-order CFA models fitted to the full cognitive dataset (N = 1,482 observations). Unlike the primary analysis, in which the CFA was fitted separately within each training set and applied to the corresponding test set to prevent data leakage, this analysis was intended to assess the robustness of the latent construct itself rather than the prediction pipeline. Accordingly, both models were estimated using the full dataset.

To inform the second-order CFA, we first conducted an exploratory factor analysis (EFA) using the psych package (v2.5.6). Parallel analysis supported a two-factor solution. An EFA with oblimin rotation indicated that SSRT reaction time standard deviation, CPT d′ catch trials, N-back 2-back accuracy, and WIAT Word Reading loaded on a common factor, which we labelled Executive Functioning and Language, whereas Digit Span Forward and Digit Span Backward loaded on a separate Digit Span factor.

We then specified a second-order CFA in lavaan (v0.6-19), in which the Executive Functioning and Language and Digit Span factors loaded onto a higher-order g factor. For model identification, the variance of the second-order factor was fixed to 1. The model was estimated using maximum likelihood with full-information maximum likelihood (FIML) to accommodate missing data. Model fit was good (CFI = .981, TLI = .959, RMSEA = .069, 90% CI [.053, .086], SRMR = .024). The higher-order g factor loaded strongly on both first-order factors (standardized loadings = .898 and .905), and standardized item loadings on the first-order factors ranged from .599 to .804 (Figure S1).

Importantly, g scores derived from the second-order model were nearly identical to those obtained from the first-order model used in the primary analyses (r = .996, p < .0001; Figure S2). This near-perfect correspondence indicates that the derivation of g was highly robust to model specification and that the two operationalizations were functionally equivalent for the purposes of the present analyses.

**Fig. S1**

*Factor loadings for re-derived first-order, and second-order g used in CFA sensitivity analysis*

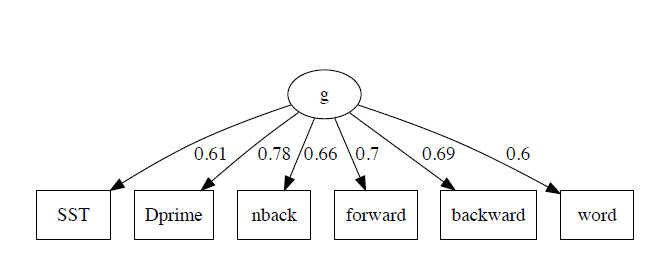

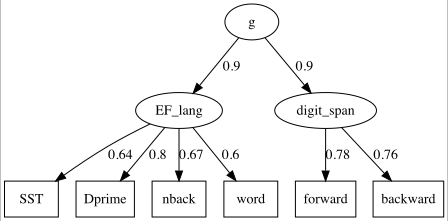

**Fig S2.**

*Scatter Plot between second-order g and first-order g*

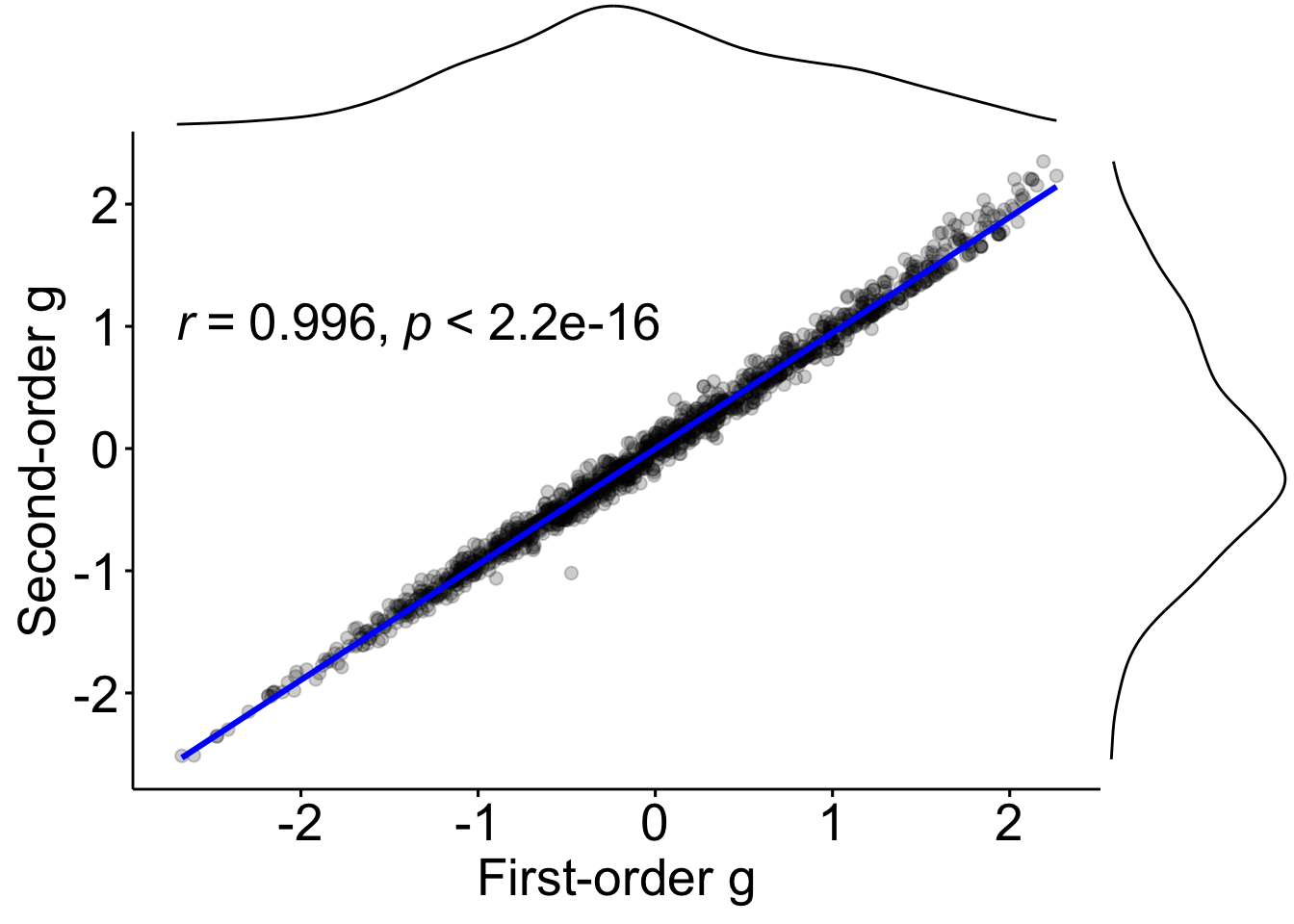

*Datapoints represent a single participant-timepoint between the two derived g-factors during sensitivity analysis. The histograms represent the data distribution of g-factors.*

### **Structural and Functional MRI data processing**

**Atlases and parcellation**
For our structural data, we utilised Destrieux cortical atlas (148 parcels, 74 per hemisphere) for cortical features^9^, and Freesurfer ASEG (19 subcortical parcels)^10^ for subcortical volume. For fMRI data, we utilised the Glasser cortical atlas^11^ (360 parcels, 180 per hemisphere) for segmentation, and 19 subcortical regions from HCP-MMP1.0^11,12^ based on FreeSurfer ASEG. For visualisation of functional connectivity, we arranged cortical vertices according to the Cole-Anticevic brain network parcellation^13^, with subcortical regions grouped into a subcortical category.

### **Neuroimaging data pre- and post- processing**

For sMRI, we preprocessed the data using fMRIPrep with the full processing pipeline. For each T1-weighted image, intensity non-uniformity was corrected using N4BiasFieldCorrection from Advanced Normalization Tools (ANTs). Brain extraction was performed using the Nipype implementation of antsBrainExtraction, with the OASIS30ANTs template as the target. Tissue segmentation into cerebrospinal fluid (CSF), white matter (WM), and gray matter (GM) was carried out using FAST (FSL v5.0.9). Cortical surface reconstruction was performed with recon-all (FreeSurfer v6.0.1), and images were spatially normalised to MNI152NLin2009cAsym standard space. The longitudinal processing option in fMRIPrep was not used, allowing structural MRI data from each time point to be processed independently and retained as separate measurements.

For fMRI, in addition to fMRIPrep, we also applied XCP-D for post-processing. XCP-D was run in LINC mode using the CIFTI input (with the default flag override). We processed functional MRI data twice, once with censoring and once without. The censored data were utilised for calculating functional connectivity (FC) and regional homogeneity (ReHo), while the uncensored data were utilised for calculating amplitude of low frequency fluctuations (ALFF).

We ran XCP-D in two streams: one for FC and ReHo and the other for ALFF. For FC and ReHo, we ran the data with censoring, where volumes with framewise displacement greater than 0.5mm were removed (fd-thresh 0.5), with minimum coverage of 0.4 (minimum proportion of non-zero, non-NaN vertices or voxels), and no minimum time (of good volumes remaining) required to process. This combination of settings was chosen so that we could then later manually assess which participants data needed to be excluded based on excessive motion and paucity of scan-time remaining. For ALFF, we disabled motion censoring (fd-thresh 0), to preserve the temporal structure necessary for ALFF calculation from timeseries. Across both of these streams, we disabled spatial smoothing, and utilised despiking of large spikes, a low-pass filter of 0.08Hz, and aCompCor^14^ for nuisance regression.

Following processing in XCP-D, we extracted motion, run, and post-censoring-remaining volume data for each subject-timepoint from quality control reports generated by XCP-D. From this data, we systematically excluded any subject-timepoint scan run from censored data with less than 50% of original volumes remaining following censoring, and summarily excluded any subject-timepoint with less than two remaining good runs (representing 10-minutes of scan time), or less than 50% of total volumes remaining across all runs following censoring. We then removed those same subject-timepoints from the uncensored data. The 0.5 mm framewise displacement threshold was adopted in line with established scrubbing criteria^15^.

**sMRI and fMRI feature processing**

Four structural and three functional feature sets were extracted from MRI data. Structural features (cortical surface area, cortical thickness, subcortical volume, and total brain volume) were derived from fMRIPrep output using nilearn (0.11.1)^16^, with one observation per subject-timepoint. Functional features of ALFF and ReHo were extracted as parcellated regional means from XCP-D output files. For functional connectivity, Pearson correlations were computed between the censored time series of each pair of the 377 parcellated regions (358 cortical from the Glasser atlas, 19 subcortical from HCP-MMP1.0), Fisher Z-transformed, and where multiple scan runs were available, averaged element-wise in Fisher Z space prior to analysis.

### **Hyperparameter grids for the first stage models**

Hyperparameter grids were typically determined by the number of features in each modality, with broader search spaces applied to low and mid-dimensional feature sets and more constrained grids applied to very high-dimensional feature sets. Where modality-specific properties (e.g., tendencies toward overfitting, low feature count, or numerical instability) required departing from these general rules, modality-specific grids were applied to ensure convergence on an appropriate solution. These cases are identified in the table notes below.

**Table S5.** *Kernel Ridge Regression hyperparameter search spaces by modality and kernel type.*

| **Modality** | **Kernel** | **Alpha search space** | **Gamma search space** | **Degree** |
| --- | --- | --- | --- | --- |
| **Functional Connectivity** | Linear | 100, 200, 300, 400, 500, 600, 700, 800, 900, 1000, 5000, 10000, 20000 | N/A | N/A |
|  | RBF | 1, 10, 100, 1000, 5000, 10000, 50000 | 1e-7, 1e-6, 1e-5, 1e-4, 1e-3 | N/A |
| **ALFF** | Linear | 0.0001, 0.001, 0.01, 0.1, 0.5, 1, 5, 10, 50, 100, 200, 300, 500 | N/A | N/A |
|  | RBF | 0.0001, 0.001, 0.01, 0.1, 0.5, 1, 5, 10, 50, 100 | 1e-5, 1e-4, 1e-3, 1e-2, 0.1, 1.0 | N/A |
|  | Poly | 0.0001, 0.001, 0.01, 0.1, 1, 10, 100, 200, 300, 500 | 1e-4, 1e-3, 1e-2, 0.1, 1.0 | 2, 3 |
| **Cortical Area** | Linear | 1000, 2000, 5000, 10000, 12000, 14000, 15000, 20000, 25000, 30000, 35000, 40000 | N/A | N/A |
|  | RBF | 10, 50, 100, 200, 300, 400, 500, 1000, 2000, 5000, 10000, 15000, 20000 | 0.1, 0.5, 1.0, 2.0, 5.0 | N/A |
| **Cortical Thickness** | Linear | 0.0001, 0.001, 0.01, 0.1, 0.5, 1, 5, 10, 50, 100, 200, 300, 500 | N/A | N/A |
|  | RBF | 0.0001, 0.001, 0.01, 0.1, 0.5, 1, 5, 10, 50, 100 | 1e-5, 1e-4, 1e-3, 1e-2, 0.1, 1.0 | N/A |
|  | Poly | 0.0001, 0.001, 0.01, 0.1, 1, 10, 100, 200, 300, 500 | 1e-4, 1e-3, 1e-2, 0.1, 1.0 | 2, 3 |
| **ReHo** | Linear | 0.1, 0.5, 1, 5, 10, 50, 100, 500, 1000 | N/A | N/A |
|  | RBF | 0.1, 0.5, 1, 5, 10, 50, 100, 500, 1000 | 1e-4, 1e-3, 1e-2, 0.1 | N/A |
|  | Poly | 0.1, 0.5, 1, 5, 10, 50, 100 | 1e-3, 1e-2, 0.1 | 2, 3 |
| **Subcortical Volume** | Linear | 100, 200, 300, 400, 500, 1000, 1500, 2000, 2500, 3000, 3500, 4000, 5000, 10000, 20000 | N/A | N/A |
|  | RBF | 100, 500, 1000, 5000, 10000 | 0.01, 0.1, 0.5, 1.0, 2.0 | N/A |
|  | Poly | 100, 200, 300, 400, 500, 600, 700, 800, 900, 1000, 1500, 2000, 2500, 5000 | 0.01, 0.1, 0.5, 1.0 | 2, 3 |
| **Total Brain Volume** | Linear | 100, 200, 300, 400, 500, 1000, 1500, 2000, 2500, 3000, 3500, 4000, 5000, 10000, 20000 | N/A | N/A |
|  | RBF | 100, 500, 1000, 5000, 10000 | 0.01, 0.1, 0.5, 1.0, 2.0 | N/A |
|  | Poly | 100, 200, 300, 400, 500, 600, 700, 800, 900, 1000, 1500, 2000, 2500, 5000 | 0.01, 0.1, 0.5, 1.0 | 2, 3 |

*Note. Grids for Cortical Area, ReHo, Subcortical Volume, and Total Brain Volume were modality-specific named cases. ALFF and Cortical Thickness received the general lower-dimensional grid (feature count < 1000). Functional Connectivity received its own named case reflecting its extreme dimensionality (70,876 features) and predominantly linear signal structure. The polynomial kernel was excluded from the Cortical Area grid due to severe overfitting observed during development. Gamma = N/A for linear kernel (no kernel bandwidth parameter). Degree = N/A where polynomial kernel was not included.*

**Table S6.** *XGBoost hyperparameter search spaces by modality.*

| **Modality** | **eta** | **max_depth** | **min_child_weight** | **subsample** | **colsample_bytree** | **gamma** | **reg_alpha** | **reg_lambda** | **Additional parameters** |
| --- | --- | --- | --- | --- | --- | --- | --- | --- | --- |
| **Functional Connectivity** | 0.01 | 2 | 20 | 0.5 | 0.3, 0.5 | 0.5, 1 | 1, 2 | 3, 5 | colsample_bylevel: 0.2, 0.3; max_bin: 128, 256; n_estimators: 1000, 1500 |
| **ALFF** | 0.01, 0.05, 0.1 | 3, 4, 5 | 3, 5 | 0.7, 0.8, 0.9 | 0.5, 0.7 | 0.1, 0.3 | 0, 0.5 | 1, 2 | - |
| **Cortical Area** | 0.01, 0.03, 0.05 | 2, 3, 4 | 5, 10, 15 | 0.6, 0.7, 0.8 | 0.5, 0.6, 0.7 | 0.1, 0.5, 1.0 | 0, 0.5, 1, 5, 10, 20 | 1, 2, 3, 4, 5, 10, 20, 50 | - |
| **Cortical Thickness** | 0.01, 0.05, 0.1 | 3, 4, 5 | 3, 5 | 0.7, 0.8, 0.9 | 0.5, 0.7 | 0.1, 0.3 | 0, 0.5 | 1, 2 | - |
| **ReHo** | 0.01, 0.05, 0.1 | 3, 4, 5 | 3, 5 | 0.7, 0.8, 0.9 | 0.5, 0.7 | 0.1, 0.3 | 0, 0.5 | 1, 2 | - |
| **Subcortical Volume** | 0.03, 0.05, 0.1 | 4, 5, 6 | 1, 3 | 0.9, 1.0 | 1.0 | 0 | 0 | 1 | - |
| **Total Brain Volume** | 0.03, 0.05, 0.1 | 4, 5, 6 | 1, 3 | 0.9, 1.0 | 1.0 | 0 | 0 | 1 | - |

*Note. Fixed parameters applied across all XGBoost models: booster = gbtree; objective = reg:squarederror; tree_method = hist; random_state = fixed seed. Cortical Area received a named special case grid with substantially stronger regularisation (reg_alpha, reg_lambda) reflecting observed overfitting tendencies. Functional Connectivity received the extreme-dimensionality grid (> 30,000 features), which additionally searched over colsample_bylevel, max_bin, and n_estimators. Subcortical Volume and Total Brain Volume received the ultra-low-dimensional grid (feature count <= 20), permitting deeper trees and minimal subsampling. ALFF, Cortical Thickness, and ReHo all received the medium-dimensional grid (101-5,000 features).*

**Partial Least Squares: adaptive component search**

For Partial Least Squares regression, the hyperparameter search space consisted of the number of latent components (n components), searched as integers from 1 to an adaptively determined maximum. The maximum number of components was first capped at 20 (the default ceiling), then further constrained to not exceed the number of available features or n samples - 1, whichever was smaller. For high-dimensional modalities (feature count > 10,000), the ceiling was additionally reduced to 10 components to limit overfitting; for moderately high-dimensional modalities (feature count > 1,000), the ceiling was reduced to 15 components. The optimal number of components was selected via 5-fold cross-validation within the training set of each outer fold, using mean squared error as the selection criterion.

### **Feature importance for functional connectivity**

To characterise which functional networks contributed most to stacked model predictions, we computed Pearson correlations between each Fisher Z-transformed functional connectivity value and the stacked model's predicted g-scores, pooled across all outer-fold test sets. For each of the 70,876 unique pairwise connections, the Fisher Z-transformed connectivity value was extracted across all matched subject-timepoints in the test sets, and its Pearson correlation with the corresponding stacked model predicted g-score was computed. To derive network-level summaries, connections were aggregated by Cole-Anticevic network assignment. For each network, all connections where either endpoint region belonged to that network were included. Connection-level *r* values were Fisher Z-transformed, averaged within each network, and back-transformed to r to yield a signed mean correlation (mean *r*). A mean absolute correlation (mean |r|) was derived identically but applied to the absolute values of connection-level *r* prior to averaging, providing a sign-agnostic index of overall association magnitude. Networks were ranked by the absolute value of the signed mean correlation. Both mean *r* and mean |*r*| are reported in Table S7.

**Table S7.** *Network-level correlations between predicted g-scores and functional connectivity, ranked by mean Pearson r. Mean r indicates the direction of the relationship; Mean |r| indicates the absolute magnitude. N connections refers to the number of individual connections where at least one node originated within a network.*

| Rank | Network | Mean r | Mean \|r\| | N connections |
| --- | --- | --- | --- | --- |
| 1 | Frontoparietal | -0.140 | 0.147 | 17,575 |
| 2 | Cingulo-Opercular | -0.127 | 0.134 | 19,516 |
| 3 | Default | -0.118 | 0.129 | 26,026 |
| 4 | Dorsal Attention | -0.115 | 0.122 | 8,395 |
| 5 | Somatomotor | -0.110 | 0.123 | 13,923 |
| 6 | Subcortical | -0.107 | 0.115 | 6,256 |
| 7 | Orbito-Affective | -0.101 | 0.105 | 2,241 |
| 8 | Visual 1 | -0.097 | 0.101 | 2,241 |
| 9 | Language | -0.096 | 0.107 | 8,395 |
| 10 | Auditory | -0.095 | 0.105 | 5,535 |
| 11 | Visual 2 | -0.090 | 0.100 | 18,873 |
| 12 | Posterior Multimodal | -0.089 | 0.104 | 2,611 |
| 13 | Ventral Multimodal | -0.038 | 0.063 | 1,498 |

*Note: Negative mean r values indicate that lower connectivity within and between network regions was associated with higher predicted cognitive functioning (g). The Ventral Multimodal network showed substantially weaker associations than all other networks.*

### **Commonality analyses full tables**

**Table S8.** *Full commonality analysis coefficients for the neuroimaging-and-age model. Values are expressed as a percentage of the total variance in observed g-scores. Negative components were set to zero prior to reporting. Key values reported in the main text (interindividual age: 3.58% unique + 9.45% shared; intraindividual age: 3.33% unique + 5.18% shared) are recoverable from this table.*

| Component type | Variance component | % total variance |
| --- | --- | --- |
| Unique | Unique to intraindividual neuroimaging (ĝ deviation) | 0.00% |
| Unique | Unique to interindividual neuroimaging (ĝ mean) | 8.72% |
| Unique | Unique to intraindividual age (deviation) | 3.33% |
| Unique | Unique to interindividual age (mean) | 3.58% |
| Pairwise shared | Shared: ĝ deviation and ĝ mean | 0.00% |
| Pairwise shared | Shared: ĝ deviation and age deviation | 5.18% |
| Pairwise shared | Shared: ĝ mean and age deviation | 0.00% |
| Pairwise shared | Shared: ĝ deviation and age mean | 0.00% |
| Pairwise shared | Shared: ĝ mean and age mean | 9.45% |
| Pairwise shared | Shared: age deviation and age mean | 0.00% |
| Three-way shared | Shared: ĝ deviation, ĝ mean, and age deviation | 0.00% |
| Three-way shared | Shared: ĝ deviation, ĝ mean, and age mean | 0.00% |
| Three-way shared | Shared: ĝ deviation, age deviation, and age mean | 0.00% |
| Three-way shared | Shared: ĝ mean, age deviation, and age mean | 0.00% |
| Four-way shared | Shared: ĝ deviation, ĝ mean, age deviation, and age mean | 0.00% |
| Total R² |  | **30.20%** |

*Note. ĝ deviation = intraindividual component of predicted g-scores; ĝ mean = interindividual component. Age deviation and age mean are the corresponding intra- and interindividual components of age. Total R² = 30.20% of total variance.*

**Table S9.** *Full commonality analysis coefficients (as % of total variance in g-scores) for the neuroimaging-and-hyperactivity model. Values are expressed as percentage of total variance in observed g-scores. Negative components were set to zero prior to reporting. Components contributing less than 0.15% are reported here but excluded from Figure 6C in the main text.*

| Component type | Variance component | % of total R² |
| --- | --- | --- |
| Unique | Unique to intraindividual neuroimaging (ĝ deviation) | 3.89% |
| Unique | Unique to intraindividual neuroimaging (g mean) | 13.21% |
| Unique | Unique to interindividual hyperactivity (deviation) | 0.00% |
| Unique | Unique to intraindividual hyperactivity (mean) | 4.48% |
| Pairwise shared | Shared: ĝ deviation and ĝ mean | 0.06% |
| Pairwise shared | Shared: ĝ deviation and Hyperactivity deviation | 1.25% |
| Pairwise shared | Shared: ĝ mean and Hyperactivity deviation | 1.71% |
| Pairwise shared | Shared: Hyperactivity mean and ĝ deviation | 0.03% |
| Pairwise shared | Shared: ĝ mean and Hyperactivity mean | 3.43% |
| Pairwise shared | Shared: Hyperactivity deviation and Hyperactivity mean | 0.00% |
| Three-way shared | Shared: ĝ mean, ĝ deviation, and Hyperactivity deviation | 0.00% |
| Three-way shared | Shared: ĝ mean, Hyperactivity mean, and ĝ deviation | 0.00% |
| Three-way shared | Shared: Hyperactivity deviation, Hyperactivity mean, and ĝ deviation | 0.01% |
| Three-way shared | Shared: ĝ mean, Hyperactivity deviation, and Hyperactivity mean | 0.00% |
| Four-way shared | Shared: ĝ mean, ĝ deviation, Hyperactivity deviation, and Hyperactivity mean | 0.04% |
| Total R² |  | 25.28% |

*Total R² = 25.28% of total variance. Negative commonality components were set to zero prior to reporting but were not renormalised; as a result, the sum of displayed components (28.08%) exceeds the total R². Individual component values are consistent with those reported in the main text.*

**Table S10.** *Full commonality analysis coefficients for the neuroimaging-and-inattention model. Values are expressed as percentage of total variance in observed g-scores. Negative components were set to zero prior to reporting. Components below 0.15% are excluded from Figure 6F in the main text but reported here.*

| Component type | Variance component | % total variance |
| --- | --- | --- |
| Unique | Unique to intraindividual neuroimaging (ĝ deviation) | 3.70% |
| Unique | Unique to interindividual neuroimaging (ĝ mean) | 12.86% |
| Unique | Unique to intraindividual Inattention (deviation) | 0.00% |
| Unique | Unique to interindividual Inattention (Inattention mean) | 11.76% |
| Pairwise shared | Shared: ĝ deviation and ĝ mean | 0.03% |
| Pairwise shared | Shared: ĝ deviation and Inattention deviation | 1.39% |
| Pairwise shared | Shared: ĝ mean and Inattention deviation | 1.73% |
| Pairwise shared | Shared: ĝ deviation and Inattention mean | 0.36% |
| Pairwise shared | Shared: ĝ mean and Inattention mean | 3.77% |
| Pairwise shared | Shared: Inattention deviation and Inattention mean | 0.00% |
| Three-way shared | Shared: ĝdeviation, ĝmean, and Inattention deviation | 0.00% |
| Three-way shared | Shared: ĝdeviation, ĝmean, and Inattention mean | 0.00% |
| Three-way shared | Shared: ĝdeviation, Inattention deviation, and Inattention mean | 0.00% |
| Three-way shared | Shared: ĝmean, Inattention deviation, and Inattention mean | 0.00% |
| Four-way shared | Shared: ĝ deviation, ĝ mean, Inattention deviation, and Inattention mean | 0.00% |
| Total R² |  | **32.55%** |

*Total R² = 32.55% of total variance. Negative commonality components were set to zero prior to reporting but were not renormalised; as a result, the sum of displayed components (35.60%) exceeds the total R². Individual component values are consistent with those reported in the main text.*

**Example of commonality analysis calculation**

interindividual age-related variance refers to the total effect of age_mean_. or the variance in observed cognitive functioning (g_observed_) explained by the between-person component of age (age_mean_) when it is entered as the sole fixed effect in the model. This is quantified by the marginal R² of the model:

g_observed_ ~ age_mean_ + (1 | Participant ID)

which equals 13.03%.

Similarly, when the between-person component of the neuroimaging marker (ĝmean) is entered as the sole fixed effect:

g_observed_ ~ ĝ_mean_ + (1 | Participant ID)

the marginal R² is 18.17%, which we refer to as the total effect of ĝ_mean_.

When both predictors are included simultaneously:

g_observed_ ~ age_mean_ + ĝ_mean_ + (1 | Participant ID)

the marginal R² increases to 21.75%. Importantly, this value is smaller than the sum of the two total effects (13.03% + 18.17% = 31.17%), indicating that part of the explained variance is shared by age_mean_ and ĝ_mean_.

This shared variance, or common effect, is calculated using standard commonality analysis principles:

Common effect = Total effect of age_mean_ + Total effect of ĝ_mean_ − Joint effect of age_mean_ and ĝ_mean_

= 13.03% + 18.17% − 21.75% = 9.45%.

Thus, of the 13.03% of variance attributable to the interindividual component of age, 9.45% is also accounted for by the neuroimaging marker. Consequently, the proportion of interindividual age-related variance captured by ĝ_mean_ is:

9.45% / 13.03% × 100 = 72.52%.

In other words, approximately 72% of the between-person age-related variance in cognitive functioning was captured by the between-person neuroimaging marker.

### **Confound sensitivity analyses**

**Methods**

To assess the robustness of the principal findings to potential confounding factors, we conducted a series of sensitivity analyses examining the influence of sex, fMRI signal quality, sMRI signal quality, stimulant medication use, and sociodemographic status.

Potential influences of sex differences on predictive performance of the stacked model were examined by comparing Pearson correlations between observed and predicted g scores across test-set observations for male (n = 634) and female (n = 406) participants. To test both for a difference and for equivalence, we conducted a conventional test for the difference between two independent correlations together with a two one-sided tests (TOST) procedure for their equivalence, applied to Fisher z-transformed correlations ^17,18^. In the absence of an a priori benchmark for a substantively meaningful sex difference, the equivalence bounds were justified using the sample-size (resource-based) approach described by Lakens et al. (2018) ^17^: the bound was set to the smallest correlation difference the available group sizes were adequately powered to detect. Given the observed group sizes, the design had 80% power to detect a Fisher z difference of 0.18 (two-sided alpha = .05), and equivalence bounds of plus or minus 0.18 Fisher z units were adopted accordingly. Statistical equivalence was concluded when the larger of the two one-sided p values fell below .05.

For other four potential confounds, we performed a residualisation analysis in which each confound, as well as all four confounds combined, was regressed out of the stacked model’s predicted g-scores via OLS regression. Residuals of the predicted g-scores were z-standardised, and within- and between-person components were recomputed via participant-level mean and deviation, as in the primary analyses. Participants with missing confound data were excluded from the relevant residualised analysis. Longitudinal mixed-effects models, variance decomposition, and commonality analyses were then re-run on the residualised predictions using the same model specifications as the primary analyses.

We operationalised fMRI signal quality as the number of volumes remaining after motion censoring. Volumes remaining post-censoring reflect the proportion of fMRI volumes retained following framewise displacement-based censoring (FD < 0.5mm). Participants with fewer usable volumes may produce less stable functional connectivity estimates. Volumes were extracted from XCP-D quality control reports and matched to each participant-timepoint.

We operationalised sMRI signal quality as the mean Euler number. The mean Euler number was derived from FreeSurfer's surface reconstruction of T1-weighted structural MRI, calculated as the mean of the left- and right-hemisphere Euler numbers, and serves as an index of structural image quality^19^. Lower (more negative) Euler numbers indicate poorer surface reconstruction quality, which may relate to head motion or image artifacts.

We operationalised current stimulant medication status as a binary indicator (1 = prescribed stimulant medication since the previous visit and/or currently taking stimulant medication; 0 = no stimulant prescription). Medication information was derived from parent-reported treatment status collected during Oregon ADHD 1000 study interviews and linked to each neuroimaging timepoint using a six-month matching window. Among participants prescribed stimulant medication, 36.4% received amphetamine-based preparations, 51.3% received methylphenidate-based preparations, and 14.3% received other stimulant medications that were not further specified.

Socioeconomic status was operationalised using parental educational attainment, as recent research suggests that neuroimaging-based prediction of cognitive performance may partly capture socioeconomic gradients^20^. Specifically, parental education was defined as the higher of maternal and paternal educational attainment and represented as an ordinal variable.

Medication status, volumes remaining, and Euler number had complete longitudinal data for 697 observations; parental education was constrained to 682 longitudinal observations (due to missingness), and the combined four-confound analysis thus yielded 682 from 268 participants. All analyses were conducted in R (4.6.0) using lme4, r2mlm, and glmm.hp libraries.

**Confound sensitivity analysis results**

For sex, Pearson correlations between observed and predicted g scores of the stacked model were compared across test-set observations for male and female participants. The two correlations were not statistically different (male r = .438, female r = .491; z = -1.04, p = .30) and were statistically equivalent within the specified bounds (Fisher z difference = -0.07; equivalence bounds plus or minus 0.18; p_equiv = .040), indicating comparable predictive performance across sexes.

Overall performance following residualisation is shown in Table S11. For fMRI, residualisation for the number of volumes remaining after censoring reduced inter-individual prediction (*r* = .35 vs. .47 before residualisation) but had minimal effect on intra-individual prediction (*r* = .52 both before and after residualisation). This may be reflected in the fact that the mean percentage of volumes remaining following censoring was 86.73% (SD = 20.55%). Thus, fMRI retained its capacity to predict within-person changes in cognitive functioning after adjustment for data quality.

For sMRI, residualising for Euler number had minimal impact on predictive performance, with largely unchanged correlations for both inter-individual prediction (*r* = .49 vs. .47 before residualisation) and intra-individual prediction (*r* = .53 vs. .52 before residualisation). These findings indicate that sMRI retained its capacity to predict both between-person differences and within-person changes in cognitive functioning after adjustment for image quality.

Likewise, residualising for stimulant medication status had negligible effects on predictive performance, yielding similar correlations for inter-individual prediction (*r* = .49 vs. .47 before residualisation) and intra-individual prediction (*r* = .54 vs. .52 before residualisation). Residualising for socioeconomic status, indexed by parental education, also produced minimal changes in performance (inter-individual: *r* = .48 vs. .47 before residualisation; intra-individual: *r* = .54 vs. .52 before residualisation). Together, these results suggest that neither stimulant medication use nor socioeconomic status substantially contributed to the predictive performance observed in the present study.

Residualising all four potential confounds simultaneously produced a pattern of results similar to that observed when residualising for the number of fMRI volumes remaining after censoring. Specifically, inter-individual prediction was attenuated (*r* = .37 vs. .47 before residualisation), whereas intra-individual prediction was largely unaffected (*r* = .52 vs. .50 before residualisation). This pattern suggests that fMRI signal quality was the primary contributor to the reduction in between-person predictive performance, with the other confounds exerting comparatively little influence. Importantly, the ability of the s/fMRI stacked marker to predict within-person changes in cognitive functioning remained robust after adjustment for all assessed confounds.

**Table S11.**

Comparison of key performance metrics across the primary analyses after residualisation

| Metric | Original | Resid-fMRI | Resid-sMRI | Resid-  med | Resid-SES | Resid-all |
| --- | --- | --- | --- | --- | --- | --- |
| Overall r | 0.459 | 0.38 | 0.47 | 0.47 | 0.47 | 0.40 |
| Interindividual r (means) | 0.47 | 0.35 | 0.49 | 0.49 | 0.48 | 0.37 |
| Intraindividual r (deviations) | 0.52 | 0.52 | 0.53 | 0.54 | 0.54 | 0.50 |
| $\boldsymbol{R}_{\boldsymbol{Interindividual}}^{\boldsymbol{2}\boldsymbol{(}\boldsymbol{f}\boldsymbol{2}\boldsymbol{)}}$ | 25.01 | 12.87 | 26.07 | 25.96 | 25.38 | 14.95 |
| $\boldsymbol{R}_{\boldsymbol{Intraindividual}}^{\boldsymbol{2}\boldsymbol{(}\boldsymbol{f}\boldsymbol{1}\boldsymbol{)}}$ | 18.82 | 18.05 | 19.01 | 20.04 | 20.27 | 17.17 |
| Intraindividual age-related change captured | 60.87 | 55.51 | 58.12 | 60.96 | 61.17 | 52.70 |
| Interindividual age-related change captured | 72.52 | 50.26 | 72.90 | 73.29 | 72.93 | 55.35 |
| Hyperactivity-cognition interindividual | 43.36 | 8.33 | 41.40 | 42.94 | 41.47 | 15.78 |
| Hyperactivity-cognition intraindividual | 100.00 | 100.00 | 100.00 | 100.00 | 100.00 | 100.00 |
| Hyperactivity -cognition captured total | 58.79 | 30.42 | 56.59 | 58.40 | 36.78 | 36.47 |
| Inattention-cognition interindividual | 25.99 | 2.78 | 23.38 | 25.24 | 24.75 | 2.60 |

*Resid-fMRI: volumes remaining post-censoring regressed out; Resid-sMRI: Resid-med: current stimulant medication status regressed out; Resid-SES: parental education regressed out; Resid-all: all four combined regressed out. Overall r reflects the correlation between residualised predictions and observed g across all participants with complete confound data; interindividual, intraindividual, and all LME-derived metrics are computed on the longitudinal subsample (≥2 timepoints). Interindividual r reflects the correlation between participant-mean residualised predictions and observed g . Intraindividual r reflects the correlation between participant-deviation residualised predictions and observed g. Intraindividual and interindividual R² values are derived from multilevel variance decomposition of the neuroimaging-only LME model. Capture percentages reflect the proportion of age-related or symptom-cognition variance shared with neuroimaging predictions, as derived from commonality analyses, expressed as proportions of the relevant age or symptom variance component. Intraindividual and total rows are omitted for inattention as Inattention Deviation was not a significant predictor of observed g at the intraindividual level in the primary analysis.*

20.	Marek, S. *et al.* Patterns of brain-wide associations reflect socioeconomics. *Science* **392**, eaee6213.
